# Unsupervised multi-animal tracking for quantitative ethology

**DOI:** 10.1101/2025.01.23.634625

**Authors:** Yixin Li, Xinyang Li, Qi Zhang, Yuanlong Zhang, Jiaqi Fan, Zhi Lu, Ziwei Li, Jiamin Wu, Qionghai Dai

## Abstract

Quantitative ethology necessitates accurate tracking of animal locomotion, especially for population-level analyses involving multiple individuals. However, current methods rely on laborious annotations for supervised training and have restricted performance in challenging conditions. Here we present an unsupervised deep-learning method for multi-animal tracking (UDMT) that achieves state-of-the-art performance without requiring human annotations. By synergizing a bidirectional closed-loop tracking strategy, a spatiotemporal transformer network, and three sophisticatedly designed modules for localization refining, bidirectional ID correction, and automatic parameter tuning, UDMT can track multiple animals accurately in various challenging conditions, such as crowding, occlusion, rapid motion, low contrast, and cross-species experiments. We demonstrate the versatility of UDMT on five different kinds of model animals, including mice, rats, *Drosophila*, *C. elegans*, and *Betta splendens*. Combined with a head-mounted miniaturized microscope, we illustrate the power of UDMT for neuroethological interrogations to decipher the correlations between animal locomotion and neural activity. UDMT will facilitate advancements in ethology by providing a high-performance, annotation-free, and accessible tool for multi-animal tracking.

## Introduction

Animal behavior is closely related to their internal state^1–3^ and external environment^4,5^. Quantifying animal behavior is a fundamental step in ecology, neuroscience, psychology, and various other fields^6–8^. As the most basic representation of behavior, the position of animals can reflect the locomotion of individuals and is an indispensable metric for behavioral analysis, especially for population-level studies involving multiple animals. Over the past few decades, the technology for animal tracking evolves continuously and recent advances have catalyzed a series of scientific discoveries^9,10^. Specifically, tracking insects in laboratories permits the identification of neural circuits and genes involved in visual navigation and locomotion control^11,12^. In the wild, statistics on individual and population-level animal migration allows the revelation of disruption and ecotoxicology caused by chemical pollution^13^. However, there exist enduring challenges impeding multi-animal tracking advancing towards higher accuracy, larger scale, and more complex scenarios, especially the similar appearance and frequent interactions of animals of the same species.

Growing demands in quantitative ethology have motivated concerted efforts to develop high-accuracy and generalized tracking methods. Recently, artificial intelligence has been increasingly adopted by researchers, achieving remarkable success in animal tracking^14–17^. As the mainstream methodology, supervised-learning-based algorithms can achieve good performance^18–20^, but they require substantial amounts of manual annotations for training (usually hundreds or even thousands of video frames), which is time-consuming and labor-intensive. Also, the workload for annotation increases linearly with the number of animals and behavioral diversity^16^. To reduce human intervention, semi-automatic annotation has been incorporated into animal tracking^21^, allowing users to interactively select optimal thresholds for animal segmentation through a graphical user interface (GUI). However, this strategy is not applicable to complex environments and low-contrast conditions where rigid thresholding for animal segmentation loses feasibility. Furthermore, in scenarios involving frequent animal interactions and occlusions, existing tracking methods are prone to identity switches (IDSW) due to insufficient mechanisms for correcting anomalous trajectories. This prevalent limitation results in accumulated errors that significantly degrade tracking accuracy.

Recently, unsupervised learning shows great potential to eliminate the reliance on human annotation or ground-truth labels by construct supervisory relationships directly from data, instead of resorting to external labels^22,23^. Latest research has demonstrated that unsupervised learning can perform better than supervised methods when applied to previously unseen datasets^24–27^, providing a feasible methodology for achieving higher accuracy with minimal annotation costs^28–30^. Moreover, theory and practice have shown that unsupervised learning can eliminate annotation bias inherent in supervised methods^31^, which is caused by human variability and mistakes, or by insufficient labeling diversity that fails to represent the entire dataset. Despite these expected advantages, the benefits of unsupervised learning have not been realized in multi-animal tracking so far, and comprehensive efforts are still needed to achieve high-accuracy tracking without requiring human annotation.

Here, we present an unsupervised deep-learning method for multi-animal tracking (UDMT) that outperforms existing tracking methods. UDMT does not require any human annotations for training. The only thing users need to do is to click the animals in the first frame to specify the individuals they want to track. UDMT is grounded in a bidirectional closed-loop tracking strategy that visual tracking can be conducted equivalently in both forward and backward directions. The network is trained in a completely unsupervised way by optimizing the network parameters to make the forward tracking and the backward tracking consistent. To better capture the spatiotemporal evolution of animal features more effectively, we incorporated a spatiotemporal transformer network (ST-Net) to utilize self-attention and cross-attention mechanisms for feature extraction, leading to a threefold reduction in IDSW compared with convolutional neural networks (CNNs). For identity correction, we designed a sophisticated module based on bidirectional tracking to relocate missing targets caused by crowding and occlusion, achieving a 2.7-fold improvement in tracking accuracy. We demonstrate the state-of-the-art performance of UDMT on five different kinds of model animals, including mice, rats, *Drosophila*, *C. elegans*, and *Betta splendens*. Combined with a head-mounted miniaturized microscope, we recorded the calcium transients synchronized with mouse locomotion to decipher the correlations between animal locomotion and neural activity. We have released the Python source code and a user-friendly GUI of UDMT to make it an easily accessible tool for quantitative ethology and neuroethology.

## Results

### Principle of UDMT

The principle of UDMT is illustrated in Fig. 1. The rationale for our unsupervised strategy is that a robust tracker should be effective in both forward and backward predictions^32^. Thus, a well-trained tracker can locate the target object along the temporal axis and subsequently track it against the temporal axis to return to its initial position (Fig. 1a). To establish the internal supervision for network training, we constructed a consistency loss to constrain the deviation between the forward tracking and the backward tracking (Methods). To capture more discriminative features, we integrated a spatiotemporal transformer network^33^ (ST-Net) to simultaneously extract spatial features of animal appearance and temporal correlations between successive frames (Fig. 1b). The encoder of ST-Net leverages self-attention blocks to aggregate multiple time-variant template features, endowing it with better capability to utilize temporal information when the animal posture changes over time. Cross-attention blocks in the decoder bridge the template branch and the search branch to determine the location of the animal inside the search area.

**Fig. 1.**
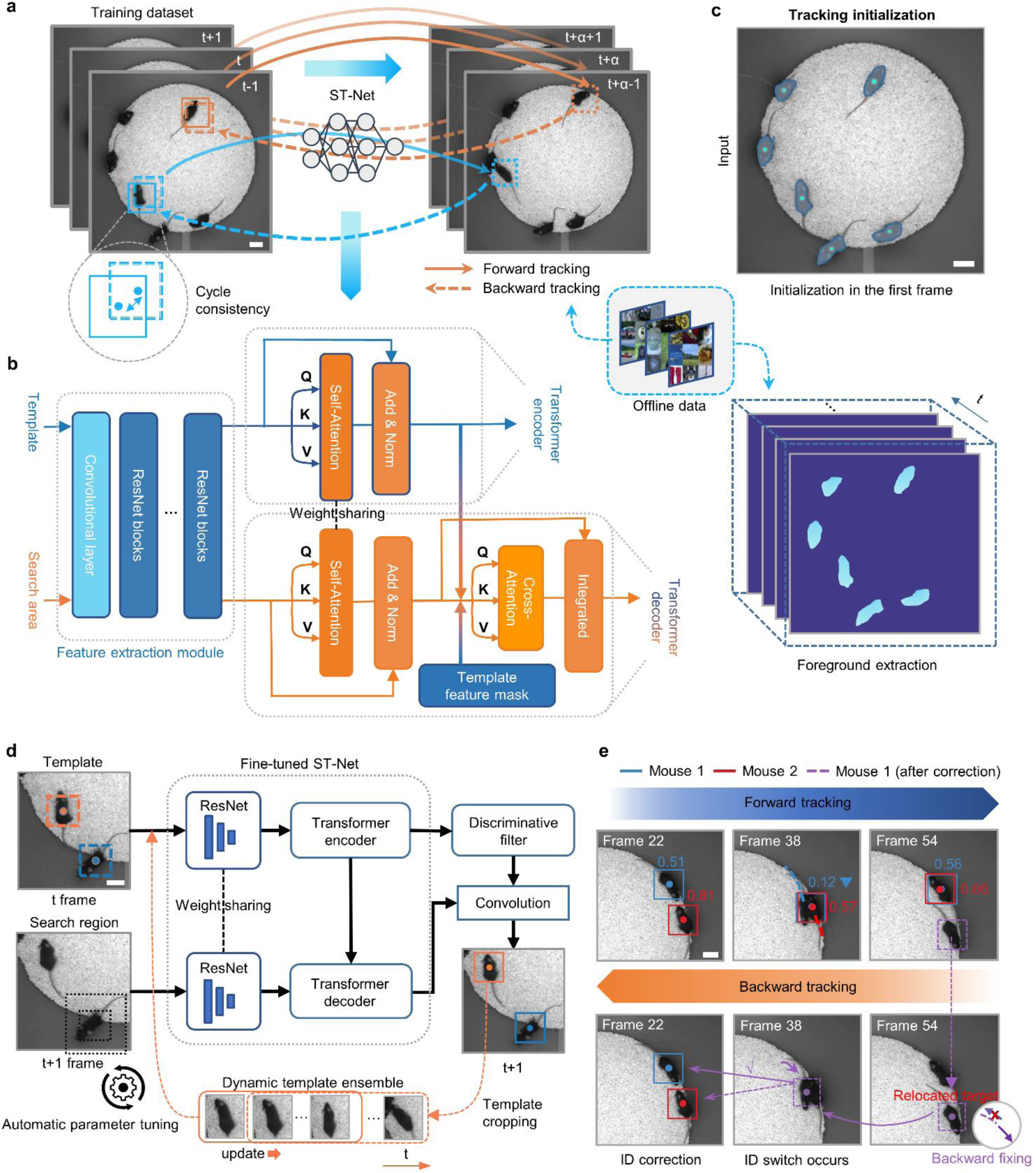
| Overview of the workflow and modules of UDMT. **a**, Self-supervised training strategy of UDMT. The video is fed into a pretrained ST-Net and the output coarse tracking results are used to construct the dataset for fine-grained training. For each animal, we track it forward and then backward to form a closed loop. The cycle-consistency loss is computed for backpropagation training of the network. **b**, The architecture of the ST-Net. It consists of a convolutional feature extraction module, a transformer encoder module, and a transformer decoder module. From left to right: the first-stage CNN takes the template and search area as input for feature extraction. These features are then passed to the following encoder and decoder. The built-in self-attention blocks leverage shared weights to convert the template and search embeddings into the same feature space. In the decoding process, cross-attention blocks are deployed to merge the template and search branch to integrate temporal contexts. **c**, Tracking initialization. Users just need to click the animal they want to track in the first frame. Then a pretrained network takes these points as input and segments animal masks throughout the whole video. **d**, Deployment of the UDMT model. The template frame (t) and search frame (t+1) are fed into the fine-tuned ST-Net simultaneously. The encoder is used to extract template features to train discriminative filters and the decoder is used to extract search features. The convolution of discriminative filters and search features can localize the object. During online tracking, the template ensemble will be continuously updated to incorporate adjacent temporal cues and adapt to the changes of target appearance. An automatic parameter tuning module is also proposed for optimal receptive field searching. **e**, Bidirectional tracking for ID correction. Those lost targets in the forward tracking process will be relocated in backward tracking, and the correct ID will be reassigned according to feature similarity. Trajectories will also be refined during backward tracking. Scale bars, 50 mm for all images.

The whole workflow contains a training process and an inference process. To initialize the training process (Fig. 1c), users just need to click the individuals they want to track using our customized GUI (Extended Data Fig. 1). Subsequently, the video and initial positions are passed to a pretained model to generate a single-object dataset for training. The pretrained model has been trained on a substantial dataset of public videos in an unsupervised manner and can achieve coarse tracking on animals. Network parameters are then optimized by stochastic gradient descent of the cycle-consistency loss. After training converges, specific representations can be learned from the dataset and memorized in specific models. To further improve tracking accuracy, we designed a localization refining module that utilizes a video object segmentation model^34^ to segment the animals of the entire video (Supplementary Fig. 1). This segmentation model is well-pretrained and does not require additional training. With our unsupervised training strategy and auxiliary operations, the original video is sufficient for training without any manual annotation.

In the inference process, search regions centered on selected animals are cropped out from input frames to eliminate the interference of redundant surrounding pixels. The cropped search regions are then fed into the trained ST-Net to infer animal positions in the next frame (Fig. 1d). Predicting the next position using multiple previous frames can utilize the continuity priors of animal locomotion. Since the posture of animals would change drastically over time in behavioral recordings, we developed a method for template updating that can continuously refresh the template ensemble throughout the tracking process to capture dynamic animal features. For deep-learning-based tracking, the size of search region is a critical hyperparameter that determines the spatial extent of the image processed by the network. However, its optimal setting is highly dependent on various factors, including movement speed, animal size, pixel size, and recording frame rate, and therefore necessitating extensive manual adjustment. In general, a larger search region provides more spatial information with the expense of introducing interference in crowded scenes, thus leading to IDSW. To obtain the optimized search region, we designed an automatic parameter tuning module based on the observation that the tracking accuracy is closely related to several explicit metrics, including the number of identity (ID) corrections, off-target localizations, and missing targets (Extended Data Fig. 2a and Supplementary Table 1). Quantitative results show that our parameter tuning module can automatically find the best search region size suitable for a specific dataset without any human intervention and lead to the highest tracking accuracy (Extended Data Fig. 2b).

Another critical challenge in multi-animal tracking is ID error owing to frequent animal interaction and occlusion. If an animal is assigned an incorrect ID, this error would propagate throughout subsequent tracking and greatly degrades tracking accuracy. To prevent cumulative ID error, we designed an ID correction module based on bidirectional tracking (Fig. 1e) that can automatically relocate missing targets and reassign correct IDs by detecting the abnormal fluctuations of moving speed and localization confidence (Extended Data Fig. 3). After ID correction, those missing and misidentified targets in forward tracking will be fixed through backward tracking.

We verified the effectiveness of UDMT on top-view behavioral recordings of multiple mice inside a circular arena (Fig. 2a and Supplementary Fig. 2a). For comprehensive comparison of each algorithm configuration, we calculated the Higher Order Tracking Accuracy (HOTA)^35^, Multi-Object Tracking Accuracy (MOTA)^36^, Identification F1-score (IDF1)^37^ and IDSW between predicted trajectories and manually annotated ground truth. To quantify the benefit of unsupervised learning, we compared the performance of the pretrained model, training from randomly initialized network parameters, and fine-tuning from the pretrained model (Fig. 2b). The result shows that unsupervised transfer learning can bring obvious improvement to tracking accuracy, especially in IDSW, which is reduced by 13-fold (1.20 ± 1.10 versus 15.60 ± 1.67, mean ± s.d.) compared to the original pretrained model and 5-fold (1.20 ± 1.10 versus 6.00 ± 0.00) compared to training from random initialization. We also conducted a progressive ablation study to assess the contributions of all proposed components (Fig. 2c), confirming that each component not only improves individual aspects of tracking but also works synergistically to achieve state-of-the-art performance. In addition to quantitative metrics, we scrutinized tracking results by visualizing trajectories (Fig. 2d), which indicates that the results of UDMT is highly consistent with the manually annotated ground truth.

**Fig. 2.**
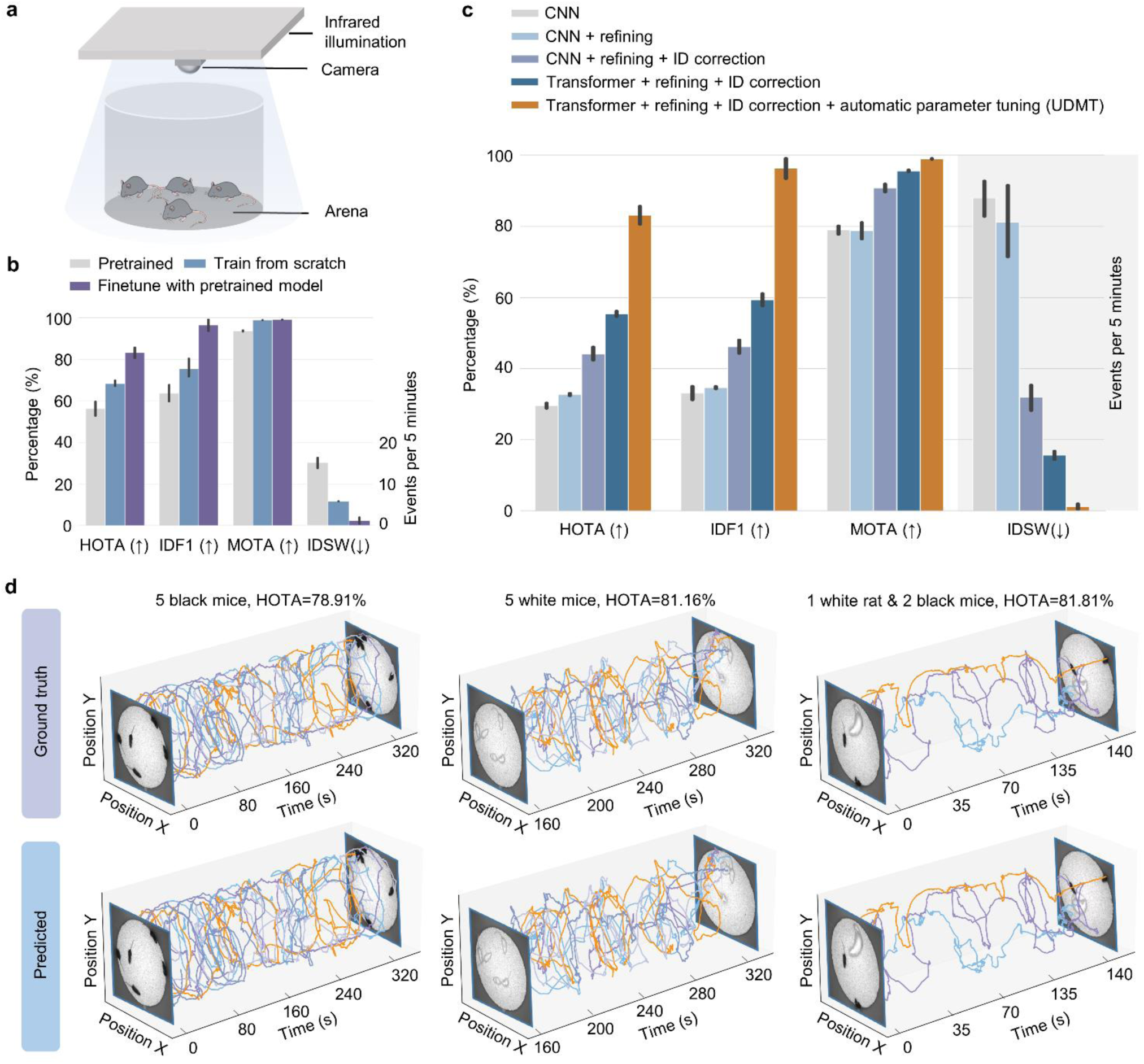
| Evaluating the performance and effectiveness of UDMT. **a**, Mouse behavioral recording system. **b**, Tracking performance (quantified by HOTA, MOTA, IDF1, and IDSW) of pretrained models, training from randomly initialized network parameters (training from scratch), and fine-tuning from the pretrained models (unsupervised transfer learning). **c**, Effectiveness of our transformer architecture and the three critical modules (localization refining, ID correction and automatic parameter tuning) in UDMT. Bars and whiskers represent mean values and 95% confidence intervals, respectively. Videos recording 7 mice (67 Hz frame rate, 29,550 frames, N=5) were used for quantitative evaluation. **d**, Manually annotated ground truth and UDMT tracking results of 5 black mice (left), 5 white mice (middle), 1 rat and 2 black mice (right). Different colors represent different individuals.

### State-of-the-art performance of UDMT in various conditions

The diversity and complexity of animal behavior pose significant challenges for tracking, including crowding, occlusion, rapid motion, and interactions. We first tested the effectiveness of UDMT in various tracking scenarios (Fig. 3a), including crowded and occluded black mice (Supplementary Video 1), low-contrast white mice (Supplementary Video 2), cross-species experiments containing 1 white rat and 2 black mice (Supplementary Video 3), and sudden rapid motion like jumping (Supplementary Video 4). With manual annotations as the ground truth, we found that UDMT can successfully handle these challenging conditions without losing or misidentifying animals. The tracking of multiple white mice also reflects that UDMT has a good tolerance for the contrast between animals and the background. Next, we performed comprehensive benchmarking against existing tracking methods to quantify the superiority of UDMT over state-of-the-art methods, including DeepLabCut (DLC)^9,18^, SLEAP^19^, and idtracker.ai (IDT.ai)^21^. We captured and compiled an ethological dataset containing 3 to 10 mice with recording frame rates ranging from 22 Hz to 94 Hz (Supplementary Table 2). For multi-animal tracking, there are three dominant factors that affect performance: the number of animals, recording duration, and frame rate. We started by comparing the accuracy of different methods for tracking different numbers of animals behaving in the same arena (Fig. 3b). Quantitative results show that UDMT outperforms other tracking methods, particularly when the number of mice is larger than five. Specifically, UDMT achieved a HOTA of 71.87 ± 3.20% when tracking 10 mice while SLEAP, IDT.ai, and DLC achieved only 49.71 ± 3.72%, 25.98 ± 2.74%, and 32.00 ± 4.85%, respectively. Such an improvement is mainly attributed to its capability to automatically detect and correct IDSW during tracking. We then evaluated the performance of different methods on different recording durations (Fig. 3c). Statistical analysis indicates that UDMT has the best accuracy for both short and long videos. For a long video over 600 s, UDMT achieved a HOTA of 71.46 ± 1.54%, much higher than other methods (IDT.ai 53.87 ± 0.61%, DLC 34.83 ± 0.69%, SLEAP 34.20 ± 1.02%), demonstrating great potential for tracking long-term ethological recordings.

**Fig. 3.**
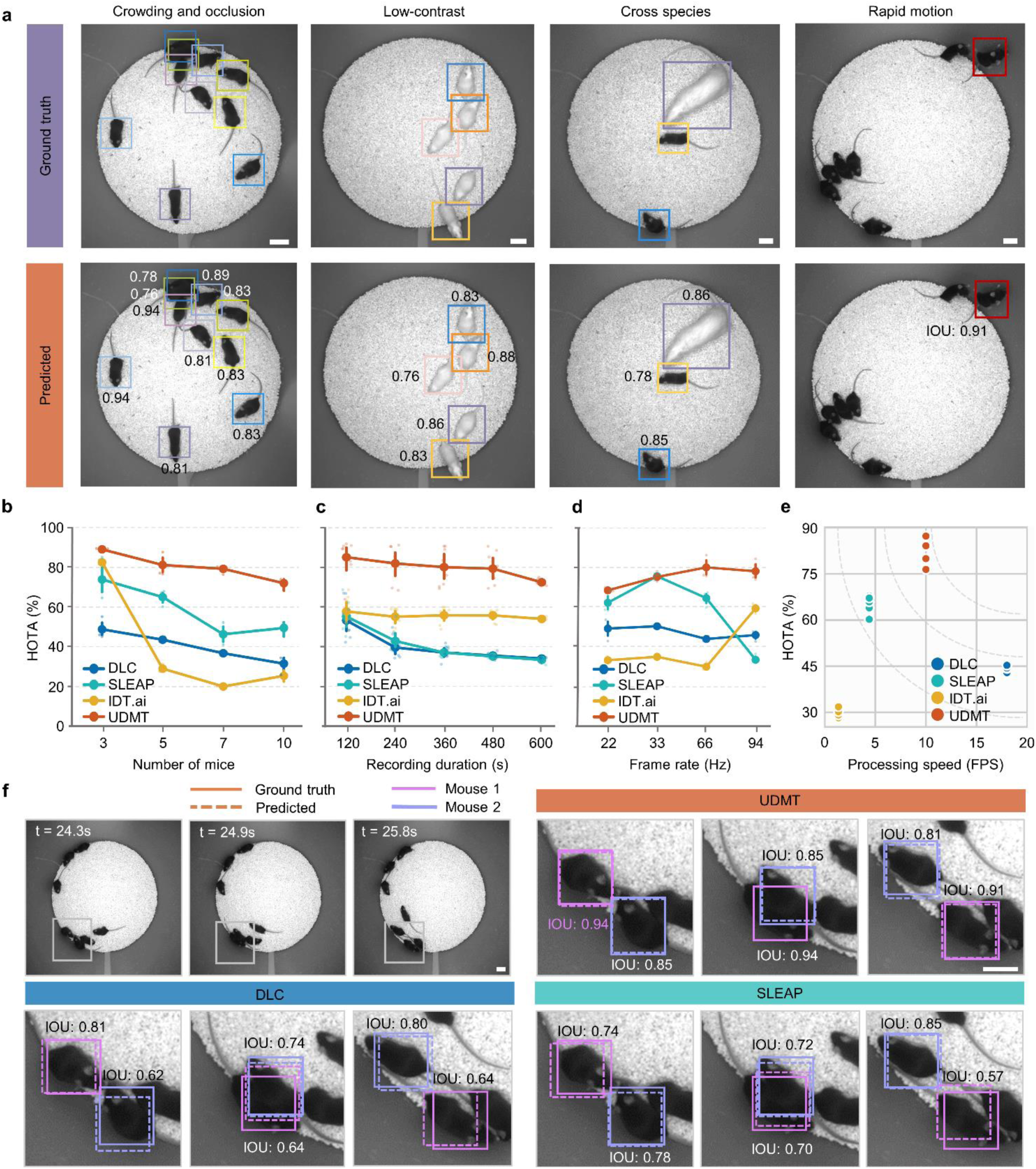
| State-of-the-art tracking performance of UDMT in various conditions. **a**, Qualitative evaluation of UDMT in various demanding recording conditions. Numbers near the boxes represent the intersection over union (IOU) between predicted and ground-truth bounding box of target animals. **b**, The relationship between the number of mice and tracking accuracy of different tracking methods. **c**, The relationship between recording duration and tracking accuracy of different tracking methods. **d**, The relationship between recording frame rate and tracking accuracy of different tracking. N=5 independent videos for all experiments. Lines represent mean values and error bars represent 98% confidence intervals. Translucent points indicate all samples. **e**, Processing speed versus accuracy of state-of-the-art tracking methods and UDMT. The 5-mouse dataset (33 Hz frame rate, 18,000 frames, N=5) was used in this experiment for quantitative evaluation. **f**, Example frames and magnified views of a 7-mouse video at three different time points. Tracking results and corresponding IOU metrics of UDMT, DLC and SLEAP are shown. Scale bars, 50 mm for all images.

Since our method relies on the similarity between two adjacent frames, it tends to be more difficult to track animals in low-frame-rate recordings. We conducted independent repeated experiments to investigate the influence of frame rate on the performance of different methods (Fig. 3d), which shows that UDMT performs better than other methods over a wide range of frame rates (from 22 Hz to 94 Hz). Another advantage of UDMT is the good balance between processing speed and tracking accuracy. It can achieve a processing speed of up to 10 frames per second and a HOTA of 80.92 ± 4.72% at the same time (Fig. 3e). We also evaluated the performance of these methods on different animal configurations, all showing that our method has the best tracking performance quantified by multiple metrics such as HOTA, MOTA, IDF1, and IDSW (Extended Data Fig. 4). Along with quantitative metrics, we also provide snapshots of the tracking results to visualize the accuracy of animal localization (Fig. 3f). Benefitting from the localization refining module, the prediction of UDMT is more consistent with the ground truth than other methods. Furthermore, we assessed how tracking performance is influenced by image quality. Once trained on a specific recording resolution or brightness, UDMT demonstrated stable superiority over other methods across a wide range of image resolution and brightness (Extended Data Fig. 5), making it a valuable tool for diverse applications. It is worth noting that our method has robust tracking performance even at resolution as low as 0.66 mm per pixel, which can be easily achieved by most ethological platforms. Additionally, UDMT is slightly affected by illumination fluctuations (Supplementary Fig. 3) but is sensitive to noise (Supplementary Fig. 4). Therefore, for data with a low signal-to-noise ratio, denoising algorithms^38–40^ should be adopted before tracking to mitigate the degradation caused by noise.

### Neuroethological analysis of multiple freely behaving mice

Deciphering how neural circuits manipulate animal behavior is a long-sought goal of contemporary neuroscience. Neural circuit interrogation of freely behaving animals with single-cell resolution is an emerging technology that promises to bring a series of breakthroughs^41–44^. In neuroethology, accurate behavioral tracking is considered as important as neural functional imaging since they jointly provide a complete chain for understanding the correlations between neural activity and animal behavior^41^. For high-accuracy neuroethological analysis of multiple freely behaving mice, we synergized UDMT with a head-mounted miniaturized microscope to investigate the influence of spatial position and velocity on neural circuits. We built a neuroethological platform to record the naturalistic behavior of multiple mice (Fig. 4a), one of which was equipped with a head-mounted miniaturized wide-field microscope weighing 2.5 g^45^. Empowered by a specially designed data processes pipeline for neuron extraction, this miniaturized microscope can record the neural activity across a 3.6×3.6 mm^2^ field of view at 4 μm lateral resolution (Methods). The collaboration between the miniaturized microscope and the top infrared camera can capture calcium transients of large-scale neural ensembles synchronized with mouse locomotion and interaction.

**Fig. 4.**
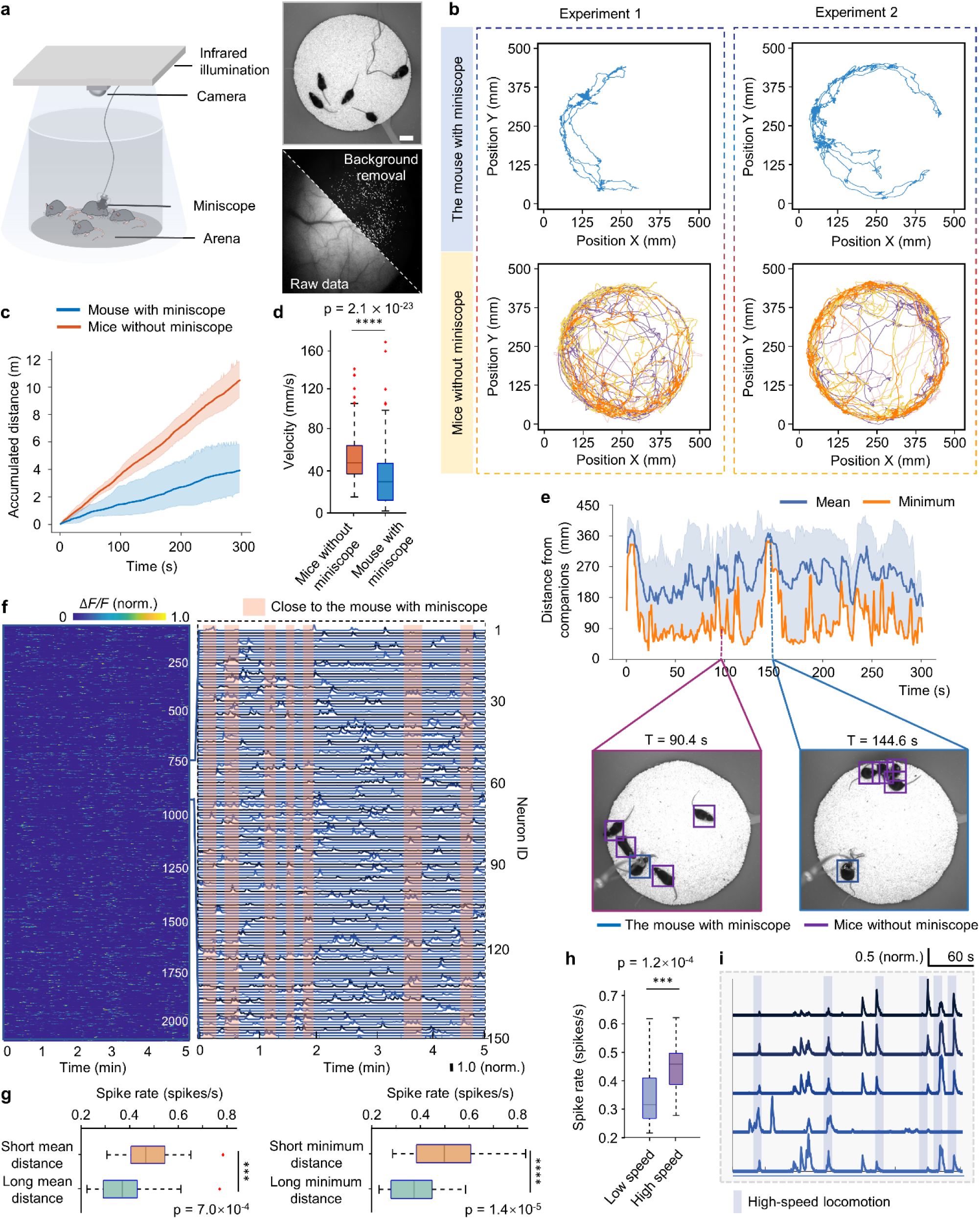
| Neuroethological analysis of multiple mice combined with a head-mounted miniaturized microscope. **a,** Neuroethological platform for simultaneous mouse behavior recording and neural calcium imaging (left). Example image of behavior recording (top right; scale bar, 50 mm) and calcium imaging (bottom right; scale bar, 500 μm, maximum intensity projection). Fluorescence background in raw calcium imaging data was removed by the processing pipeline^65^. **b**, Trajectories of the mouse with head-mounted miniaturized microscope (miniscope) and the other four mice without miniscope in two independent experiments. Different colors represent different mice. **c**, Accumulated distance over 5 minutes of moving. The orange line indicates the mean moving distance of the four free mice and the blue line indicates the moving distance of the mouse with miniscope. Shaded regions denote 95% confidence intervals. Five video (N=5) recording the movement of 5 mice were used for quantitative analysis (64 Hz frame rate, 19,380 frames per video). **c**, Accumulated distance over 5 minutes of moving. The orange line indicates the mean moving distance of the four free mice and the blue line indicates the moving distance of the mouse with miniscope. Shaded regions denote 95% confidence intervals. Five video (N=5) recording the movement of 5 mice were used for quantitative analysis (64 Hz frame rate, 19,380 frames per video). **d**, Box plot showing the speed of the mouse with miniscope and the mean speed of the other four mice without miniscope. The speed is averaged over 1 s. N=302. P values were calculated by one-sided paired t-test. ****P<0.0001. **e**, Line plot showing the distance of the mouse with miniscope to the other four mice as a function of time (top) and two representative video frames (bottom). Blue line and shaded region denote mean and range of values, respectively. **f**, Neural activity of 2,131 detected neurons in a 5-minute recording. The zoom-in panel shows the calcium traces of 200 neurons. Red shaded areas represent time windows when the distance between the four free mice and the mouse with the miniscope falls below the lower quartile of their average distances. **g**, Box plots showing neuronal spike rate at different mean and minimum spatial distances. P values were calculated by one-sided paired t-test. ***P<0.001, ****P<0.0001. **h**, Box plots showing neuronal spike rate at different moving speed. P values were calculated by one-sided paired t-test. ***P<0.001. **i**, Tuning analysis on a single-neuron level. Calcium traces of neurons that were upregulated by high-speed locomotion. Blue shadows indicate the time of high-speed locomotion.

We used 5 transgenic mice (1 equipped with the miniaturized microscope and 4 without) expressing genetically encoded GCaMP6f calcium indicators^46^ specifically in Layer 2/3 neurons. Throughout the recording duration, the miniaturized microscope can capture calcium dynamics of more than 2,000 neurons in the primary visual cortex. To associate the neural activity with mouse locomotion, we utilized UDMT to track all five mice and extract their trajectories (Fig. 4b and Supplementary Video 5). Visual inspection suggests obvious variations in the movement trajectory of the mouse equipped with the miniaturized microscope. To quantify the difference, we calculated the accumulated moving distance and instantaneous velocity of each mouse, and analyzed the difference in moving patterns between the mouse with the miniaturized microscope and the other mice without the miniaturized microscope (Fig. 4c). Further statistical result reveals that the mouse with the miniaturized microscope showed significantly reduced moving distance and velocity (Fig. 4d). Such a reduction in mobility is caused by the weight of the microscope, as well as the weight and tension of the signal transmission cables. Developing head-mounted microscopes that are lighter in weight, and reducing the weight and tension of the cables will enable more naturalistic behaviors of mice^47^.

According to our observation, we hypothesized that the aggregation status of mice would have a significant effect on neural activity. To verify our hypothesis, we visualized the distance between the mouse with the miniaturized microscope and the other four mice throughout the recording (Fig. 4e). The mean distance is the average distance from the mouse with the microscope to all other mice, and the minimum distance represents the distance from the mouse with the microscope to the nearest mouse. Specifically, when the minimum distance is large, it means that there are no other mice near the mouse with microscope. When the mean and minimum distance are very small, the mouse with microscope is likely to be surrounded by other mice. Combined with synchronized calcium imaging of 2,131 neurons over a 5-minute recording period, we quantitatively analyzed the correlations between neural activity and the relative position of the mice (Fig. 4f). We find that the overall neuronal spike rate is statistically related to the distance of the mouse from its companions. When the mouse has companions nearby, the spike rate of observed neurons is inclined to increase (Fig. 4g). In addition to distance, the first-order differential of spatial position, the velocity, is also significantly correlated to the spike rate of neurons (Fig. 4h). When the mouse undergoes rapid locomotion, its neuronal spike rate is inclined to increase. We also identified five upregulated neurons that are most sensitive to moving velocity (Fig. 4i). In brief, the tracking capability of UDMT facilitates neuroethological analysis and reveals the mechanism that neurons in the primary visual cortex tend to be more active when the mouse is surrounded by conspecifics or in high-speed locomotion.

### Tracking various model animals with UDMT

Previous experiments have verified the superiority of our method on rodents. Although rodents are the most commonly used model animals in laboratories, ethological research has been extended to various kinds of animals such as worms, insects, and fish^48–50^. To validate the broad applicability of UDMT, we applied it to other model animals such as fruit flies (*Drosophila*), worms (*Caenorhabditis elegans*), and betta fish (*Betta splendens*). The body size and behavioral patterns of these species are quite different from those of rodents. For the tracking of flies, we constructed a recording system to capture the locomotion of multiple flies inside a large culture dish (Fig. 5a,b and Supplementary Fig. 2b). We put 17 flies in the culture dish at the same time and recorded their free movement for 500 s. After extracting their movement trajectories using UDMT, we visualized all the trajectories and further computed the velocity and acceleration of these flies (Fig. 5c,d and Supplementary Video 6). The results show that the moving velocity and acceleration of these flies continued to decrease after being transferred to the arena, which is probably related to the fact that they finished exploration and gradually became familiar with the new environment. We also performed a more fine-grained analysis by visualizing the trajectories in a three-dimensional (x-y-t) manner. We identified two chasing flies with highly similar trajectories but exhibited a short temporal delay (Fig. 5e).

**Fig. 5.**
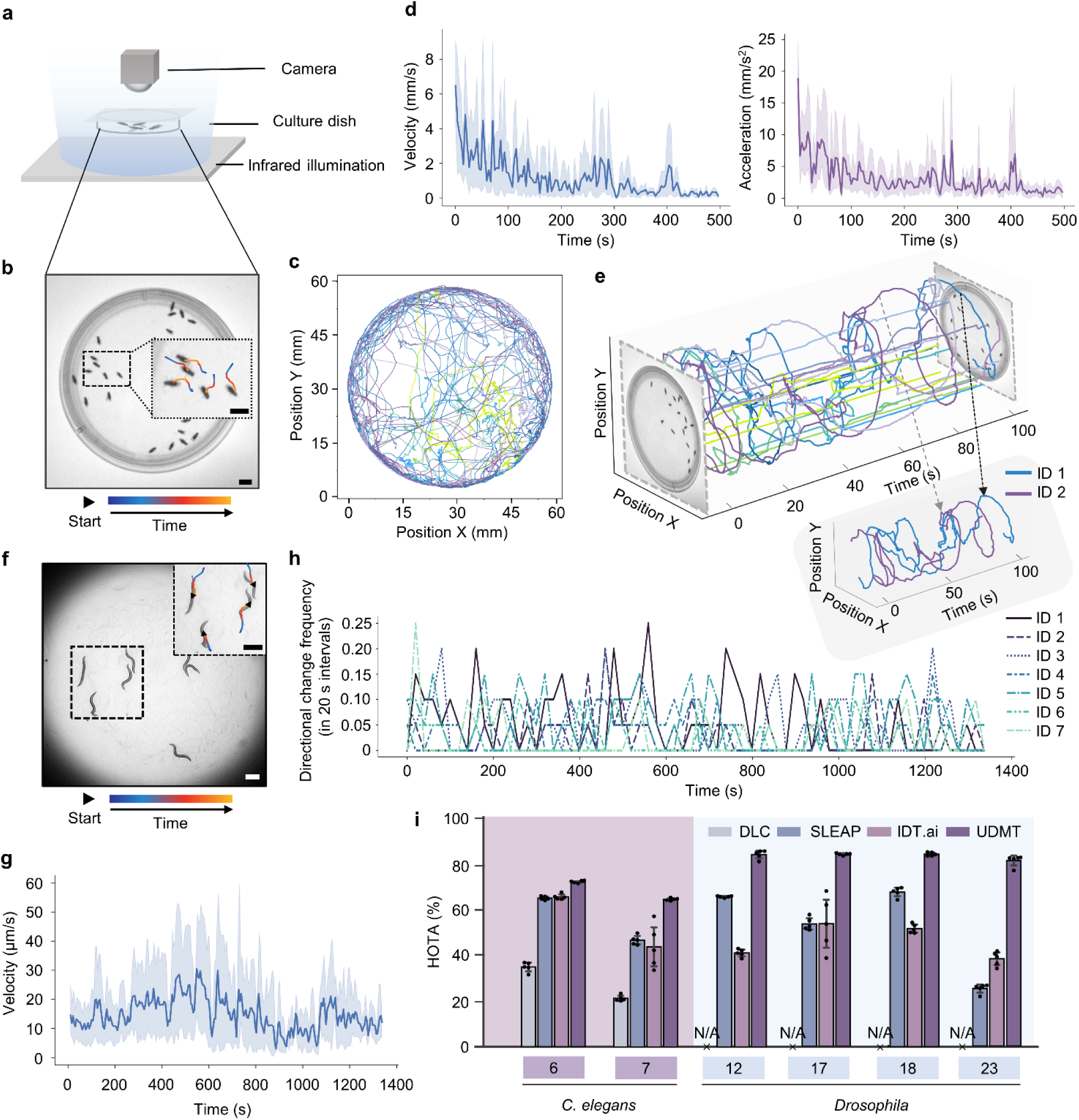
| General-purpose multi-animal tracking with UDMT. **a**, Schematic of the recording system for *Drosophila*. **b**, Example image and magnified views of the 17-*Drosophila* dataset. Scale bar, 5 mm. **c**, Projected trajectories of the 17 *Drosophila* during the entire recording period. **d**, Velocity (left) and acceleration (right) of all 17 *Drosophila* averaged over 0.2-s intervals as a function of time. Lines and shaded region denote mean and 95% confidence intervals, respectively. **e**, 3D (x-y-t) trajectories of the 17 *Drosophila* in 100 seconds. The trajectories of two chasing *Drosophila* are shown separately in the inset. The frame rate of the 17-*Drosophila* dataset is 54 Hz and the total number of frames is 26,900. **f**, Example image of the 7-*C. elegans* dataset. Scale bar, 300 μm. **g**, Velocity of all of 7 *C. elegans* averaged over 10-s intervals as a function of time. Lines and shaded region denote mean and 95% confidence intervals, respectively. The frame rate of the 7-*C. elegans* dataset is 10 Hz and the total number of frames is 13,550 frames. **h**, Directional change frequency of seven *C. elegans*. The directional change frequency was averaged over 20-s intervals. **i**. Comparing the performance of UDMT with other tracking methods on all *C. elegans* and *Drosophila* datasets. The number of animals is indicated at the bottom of each bar plot. The bars represent mean values and error whiskers represent 98% confidence intervals. N=5 for all datasets and each black point indicates a sample. N/A means that DLC fails to track *Drosophila* datasets.

Studies on *C. elegans* have shown that movement tracking can provide valuable clues to reveal their external perturbations such as mechanical and chemical interventions, as well as their internal states like metabolic health and neural function^51^. Next, we verify the performance of UDMT on tracking multiple *C. elegans*. Using a stereoscope, we imaged the free crawling of 7 *C. elegans* inside a culture dish (Fig. 5f and Supplementary Fig. 5a). We tracked all the *C. elegans* simultaneously with UDMT and visualized their trajectories and velocities throughout the 22-minute recording (Supplementary Video 7). Since both reversal (switching from moving forward to backward) and large-angle turning are important locomotion features of *C. elegans*^52^, we derived the frequency of directional change and reversal of these *C. elegans* to quantify how often they change their moving direction (Fig. 5h and Supplementary Fig. 6). Our ethological analysis indicates that these *C. elegans* behaved stably during the recording period and their directional change frequency did not show large fluctuations. Moreover, to compare UDMT with other methods for tracking flies and *C. elegans*, we conducted independent repeated tracking experiments on different numbers of flies and *C. elegans* (Fig. 5i and Extended Data Fig. 6). Comprehensive metrics, including HOTA, MOTA, IDF1, and IDSW, have demonstrated that UDMT can track flies and *C. elegans* with higher accuracy than other methods on different numbers of animals.

Lastly, we used UDMT for the ethological investigation of betta fish, a gorgeous fish exhibiting obvious aggressive behaviors^53^. When two adult betta fish are close to each other, they will manifest evident agonistic display and may chase and fight each other subsequently. Their aggressive behaviors are commonly adopted to establish dominance and defend territory, particularly during mating season^54,55^. We built an ethological recording system for betta fish and recorded the interactions of two betta fish inside the same arena (Supplementary Fig. 5b and Supplementary Video 8). The movement trajectories of the two betta fish during the 5-minute recording were extracted using UDMT (Extended Data Fig. 7a). By visualizing the trajectories in a three-dimensional coordinate, we can find a segment of trajectories related to aggressive behaviors (Extended Data Fig. 7b). We also calculated the swimming velocity of the betta fish and correlated their aggression behavior with velocity (Extended Data Fig. 7c). Combined with corresponding images, we recognized some fixed patterns of the aggressive behavior of betta fish (Extended Data Fig. 7d). The confrontation between two fish is accompanied by a drop in swimming speed to almost zero, while chasing behavior results in a rapid rise in swimming speed to seven times their average speed. During chasing, the speed of the subordinate fish is much higher than that of the dominant fish, which means that the subordinate fish will escape quickly and then the dominant fish will follow at a relatively slow speed. Such a chasing behavior is aimed at expelling enemies, rather than predation, in which the predator must pursue the prey fiercely with a comparable speed.

## Discussion

Conventional animal tracking algorithms rely heavily on substantial manually annotated training data. Preparing annotations is time-consuming and prone to errors, particularly when the annotation workload scales with the number of animals and behavioral diversity^9,19^, hindering large-scale ethological studies of animal populations. Unsupervised learning has the potential to break the reliance on manual annotations but is challenged by the similar appearance and frequent interactions of animals. We proposed an unsupervised deep-learning method that can achieve high-accuracy tracking of multiple animals without requiring any manual annotations. Our approach, UDMT, synergizes a cycle-consistency training strategy with a spatiotemporal transformer architecture, as well as three sophisticated modules including localization refining, bidirectional ID correction, and automatic parameter tuning, cooperatively enabling high-accuracy and stable multi-animal tracking. We conducted extensive experiments on five different kinds of model animals, including mice, rats, *Drosophila*, *C. elegans* and betta fish. Quantitative results demonstrate the superiority and versatility of our method. Beyond macro-level ethology, we further combined with a head-mounted miniaturized microscope for neuroethological interrogation of mouse at a single-neuron level. UDMT facilitated the discovery of the correlations between neural activity and locomotion, and assisted the identification of upregulated neurons that are closely related to moving speed.

Although we tested tracking on up to 23 animals (Fig. 5i), this is not the upper limit. Since the performance of UDMT is not only related to the number of animals, but also depends on the image resolution and frame rate, improving image resolution and frame rate could allow our method to track more animals. It is noteworthy that the generalization ability of our method is constrained when applied to different species or varying experimental settings. To obtain optimal tracking performance, we recommend training customized models for specific species and experiment conditions.

There are still several avenues to explore in the future. While our current method can simultaneously track the locomotion of multiple animals, it cannot track multiple keypoints on the animal. As the amount of annotation required for keypoint tracking is several times higher than that of position tracking, developing unsupervised methods for keypoint tracking has greater significance for quantitative ethology. Such an advancement would greatly promote fine-grained studies related to animal posture. Moreover, unsupervised learning can eliminate the reliance on task-specific annotations, thus allowing deep learning models to learn from the vast amount of unlabeled data. Inspired by recent progress in foundation models^29,56^, we envision designing a foundation tracking model pre-trained on large-scale datasets to realize generalized multi-animal and keypoint tracking, which could provide a versatile, scalable, and high-performance solution to address the limitations of current task-specific tracking methods. In terms of applications, we anticipate extending our method to the field of ecology to study the behavior of animals in the wild. Ecological systems are embedded in much more complex environments than laboratory settings, including streams, coral reefs, or forests, and are characterized by biodiversity and large-scale populations^57^. Applying our method to ecological studies such as tracking fish and bird flocks may require appropriate refinements. At the microscale, our unsupervised multi-object tracking method may play an important role in tracking cells, organelles, and particles. However, these microscopic objects have quite different properties from macroscopic objects. For example, division and apoptosis would occur during cell movement. Incorporating more specific rules and constraints is expected to improve its applicability.

## Methods

### Behavioral recording and data annotation

To evaluate how UDMT performs across species, imaging conditions, experimental conditions and other properties of behavioral recordings that may affect tracking performance, we built a collection of diverse animal datasets of behavioral recording (Supplementary Table 2). All experiments involving animals were performed in accordance with the institutional guidelines for animal welfare and have been approved by the Animal Care and Use Committee of Tsinghua University.

#### Annotation workflow

Human annotators were instructed to label in the downsampled original video, assigning an ID to each animal and marking the minimal bounding box around its body parts. We approximated the center of the minimal bounding box as the centroid of the animal, representing its position.

#### Mouse

The black-mouse dataset was used to assess tracking performance at different numbers of animals, recording frame rate, and recording durations. The white-mouse dataset was used to evaluate tracking performance in low-contrast imaging conditions where animal brightness and color are very close to the background. Male or female C57BL/6J mice were housed in a temperature- and humidity-controlled environment on a 12-hour reversed light-dark cycle, with unrestricted access to food and water. Groups of five mice were housed per cage and used for experiments at 2.5 to 4 months of age. Behavioral assays were conducted during the active (dark) phase of mice, from 12:00 PM to 5:00 PM. To ensure environmental standardization and minimize intra-cage disturbance, mice were individually housed 24 hours before behavioral testing. Age-matched animals were used across experimental groups to eliminate potential age-related confounders.

During behavioral recording, mice moved freely in an open-field arena with a diameter of 50 cm and a height of 30 cm. Videos were captured under infrared illumination (35 × 35 cm) using a Mindvision camera (MV-SUF401GM) at 40–94 Hz. This camera provides a maximum spatial resolution of 2,048 × 2,048 pixels and a frame rate of 88 Hz at full resolution. The recording frame rate can be increased by decreasing the image resolution using the control software. Infrared brightness was regulated by an analog controller (JH-AP60-2C). The custom-assembled acquisition workstation was equipped with an Intel Xeon CPU, 32 GB RAM, a Seagate BarraCuda 1-TB hard disk drive for data storage, and a NVIDIA GeForce GTX 1070Ti graphics processing unit (GPU, 8 GB memory). The raising and recording methods of rats are the same as those used for mice. For each video, human annotators labeled one frame every 300 frames, obtaining 712 labeled frames on the mice and rat datasets.

#### Mouse with head-mounted miniaturized microscope

For neural functional imaging, we used transgenic mice bred by crossing Rasgrf2-2A-dCre with Ai148 (TIT2L-GC6f-ICL-tTA2)-D strains, expressing Cre-dependent GCaMP6f genetically encoded calcium indicator. Craniotomy surgeries were performed as previously described^45^. Briefly, mice were anesthetized with 1.5% isoflurane (v/v in oxygen), and a 4.0 mm craniotomy was created using a skull drill. After removing the skull piece, a coverslip was placed over the exposed area, and a titanium headpost was affixed to the skull to stabilize the miniaturized microscope.

The miniaturized microscope, optimized for mesoscale imaging, offers a field of view (FOV) of 3.6 × 3.6 mm, a lateral resolution of 4 µm, and a depth of field of 300 µm, with a total weight of 2.5 g. Two weeks after surgery, mice were re-anesthetized and positioned in a stereotactic frame for baseplate attachment. The baseplate was manually aligned for optimal FOV centering on the sensor, then secured with dental cement and Krazy Glue. After the glue was cured, the miniaturized microscope was detached, and the mice were returned to their home cages, now ready for imaging experiments. Before each behavioral experiment, mice were lightly anesthetized (0.5–1% isoflurane in oxygen) and head-fixed to clean the cranial window and mount the miniaturized microscope. A blue LED was then activated for fluorescence excitation. Focusing was achieved by adjusting the top housing relative to the bottom housing until neurons were clear, after which the setup was locked. Once focused, mice were released into the open-field arena. Recordings were captured at 4 Hz with a resolution of 2,592 × 1,944 pixels.

#### Drosophila

*Drosophila* (wild-type Canton S) were grown on standard cornmeal media (BuzzGro from Scientiis) in 25 °C incubators with a 12/12 h light-dark cycle. The behavioral experiments setup was placed in a dedicated experimental room with controlled humidity (60%) and temperature (25 °C). The *Drosophila* were placed in a covered polystyrene petri dish for behavioral recording, which had a diameter of 54 mm and a height of 2 mm. Infrared illumination was positioned directly above the culture dish to ensure consistent lighting within the central region (Supplementary Fig. 2b). A white plastic sheet was placed below the arena to increase the contrast between *Drosophila* and the background. Videos were recorded with the same camera used for mice experiments and obtained with standard top-view recording. Black shade cloths were positioned around the camera to minimize reflections on the glass covering the arena. To transfer *Drosophila* into the arena, they were briefly anesthetized on ice before being gently placed into the setup. Recording began after they regained consciousness and normal activity. Human annotators labeled one frame every 150 frames per video, resulting in 629 labeled frames for the *Drosophila* datasets.

#### C. elegans

*C. elegans* were cultured on nematode growth medium plates seeded with OP50 *Escherichia coli* cells. The wild-type Bristol N2 strain was provided by the Caenorhabditis Genetics Center and maintained at 20 °C. To pick *C. elegans* using a worm picker (platinum wire), we slowly lowered the tip of the wire and gently swiped it along the side of the worm, then transferred the worm to a 60-mm NGM plate seeded with an OP50 lawn. For *C. elegans* imaging, a Nikon Ti2-E inverted stereoscope was utilized (Supplementary Fig. 5a). The system was equipped with a 2× objective lens (0.1 numerical aperture, 8.5 mm working distance), providing high-resolution images suitable for detailed behavioral analysis. For each video of *C. elegans*, human annotators labeled one frame every 150 frames, resulting in 196 labeled frames.

#### Betta splendens

Male *Betta splendens* used in this experiment were raised in 600 mL glass flasks at 26±2°C under a 12-hour light/12-hour dark cycle. Each fish was placed in a separate tank within a circulating aquarium system. Fish were fed twice daily and their average length was about 5.2 cm. For behavioral recordings, two male betta fish were transferred to a 20 × 30 × 30-cm transparent glass tank filled with water with a depth of 2.5 cm (Supplementary Fig. 5b). Videos were recorded with the same camera as in the mice setup. To prevent reflections on the water surface, infrared illumination was positioned beneath the fish tank, which was supported by a transparent acrylic plate to maintain approximately a distance of 30 cm from the light source. Additionally, a thick layer of frosted cellophane was placed at the bottom of the tank to ensure uniform illumination.

### Network architecture

The network architecture of UDMT retains the topology established in TrDiMP^33^, including a feature extraction module, a transformer encoder module, a transformer decoder module and a discriminative filter. We employed ResNet-50^58^ as the backbone network within the feature extraction module to derive the initial feature maps for both the template and search regions. The primary operation within the transformer encoder is self-attention, designed to enhance the features derived from a dynamically updated template ensemble. Notably, the self-attention blocks in both the encoder and decoder share weights, enabling the transformation of template and search embeddings into a unified feature space, thus facilitating subsequent cross-attention computations. We computed the cross-attention based on the search feature and template feature. The enhanced search features extracted by the spatial attention mechanism can better highlight potential target areas. Ultimately, we constructed a discriminative filter from template features and convolved it with search features to accurately localize the target.

### Training and inference

To initialize the training process, users just need to click the individuals they want to track using our customized GUI. The individuals selected by users will be automatically segmented by a generalized model^59^ and sent to a video object segmentation model^34^ for animal segmentation throughout the entire video. The masks are utilized in the localization refining module to improve tracking accuracy. During the forward tracking phase of training, we employed the raw target positions from the preceding (t-1), current (t), and subsequent (t+1) frames to crop object regions, which serve as the templates. t is randomly sampled from all frames. Frames are cropped centered on the target position and the side length is determined by the search region size. If the cropped region extends beyond the image border, it is padded with the average intensity of the entire image, which closely approximates the background intensity. The cropped regions from t+α-1, t+α, and t+α+1 frames serve as the search frames, where α is an integer randomly sampled from the range of 1 to 10. Training image patches were extracted by sampling random translations relative to the target annotations for data augmentation. The templates and search frames will be simultaneously fed into the network to locate the position of the target animal within the search frames. In the backward tracking phase, the search frames from the previous step are used as templates, and processed by the same network. The value of the loss function is computed using the initial positions and the corresponding positions of backward tracking in t-1, t and t+1 frames. Simply using the difference between ground-truth target scores and target confidence scores as the loss function biases the learning process toward negative data samples, rather than enabling the model to achieve optimal discrimination. Moreover, using naive differences cannot address the problem of data imbalance between targets and backgrounds. To address this issue, we modified the hinge loss^60^ based on the principle of support vector machines to train UDMT models in an end-to-end manner. The loss function for each label and prediction score pair is formulated by

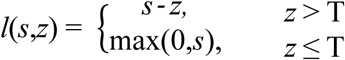

In this context, the threshold T represents the target and background regions based on the label confidence value z. For the target region where z > T, we computed the difference between the predicted confidence score s and the label z. In contrast, for the background region where z ≤ T, we only penalized positive confidence values. T was set to 0.05 in our experiment.

During the inference phase, given an annotated initial frame, subsequent frames are processed sequentially through the network. For preprocessing, each input frame is normalized by subtracting its minimum intensity value and then dividing by the difference between the maximum value and the minimum to handle the intensity variation across different videos. To leverage temporal information and adapt to changes in target appearance optimally, the template ensemble in the transformer module will be updated dynamically. Specifically, for every 5 frames, the earliest template in the ensemble is discarded, and the current template feature is appended. The feature ensemble is maintained at a maximum of 15 templates. To process the first frame, we employed data augmentation strategies (including blur, rotation, and shift) to construct an initial template ensemble containing 15 samples. Upon updating the template ensemble, we computed the new encoded template feature using our transformer encoder. While the transformer encoder is sparsely utilized (*i.e.*, every 5 frames), the transformer decoder is engaged for each frame, generating per-frame search features by propagating representations and attention cues from preceding templates to the current search patch.

All models were trained using the Adam optimizer^61^, with an exponential decay rate of 0.9 for the first moment and 0.999 for the second moment. The learning rates for the feature extraction module, transformer module, filter initialization, and filter optimization module were 0.00005, 0.001, 0.00005, and 0.0005, respectively. The ST-Net was initialized using weights provided in TrDiMP, whereas the discriminative filter was trained from scratch. The search area factor was set to 3.0, and the default number of samples (image pairs) per epoch was set to 5000. We used GPUs to accelerate the training and testing process. The batch size for all experiments was 40, which required 16.93 GB of memory per GPU when training in parallel on three GPUs. Smaller batch sizes can reduce memory demands. Training the network with a video comprising approximately 4,000 frames is sufficient to ensure satisfactory performance (Supplementary Fig. 7a). Generally, the optimal tracking performance is achieved at the 10th epochs and keeps stable within 20 epochs (Supplementary Fig. 7b). Training the model for 20 epochs on a typical dataset (with a batch size of 16) using a single Nvidia GeForce RTX 3090 GPU (24 GB memory) takes approximately 2 hours. On an already trained network, processing 16,000 frames (652 × 636 pixels, containing 5 animals) requires about 1,800 seconds. Training time can be further reduced by utilizing more powerful GPUs or parallelizing computations on multiple GPUs. PyTorch was used to construct the network and implement all operations. The tracking result of each video was saved as a separate txt file.

### Localization refining module

The localization refining module was designed to improve the accuracy of animal localization in real-time. Without the refining module, when a position deviation occurs during tracking, the error will accumulate frame by frame and degrade the accuracy of long-term tracking. For localization refining, all video frames and the initial positions of animals are sent into a pre-trained model to segment the masks of all animals. To avoid the bias of network prediction, the refining module uses the center of gravity of corresponding segment masks to replace the original localization of the network. The refined positions will be used to update subsequent tracking. The refining process is applied only for those segmentation masks whose areas are between 50% and 120% of the initial animal area, ensuring that the mask corresponds to a single animal. The initial animal area is obtained by averaging the area of all animals in the first frame. When multiple animals are in contact, the area of each animal is estimated by dividing their total area by the number of animals within that region. For some concave-shaped animals such as *C. elegans*, their centroid may not be located within segmentation masks. In these cases, the centroid of the smallest outer rectangle of each segmentation mask is used as the animal position.

### ID correction module

To reduce cumulative ID errors, we developed an ID correction module leveraging bidirectional tracking. The detailed workflow of ID correction is shown in Extended Data Fig. 3. First, if the trajectory difference index^62^ between any two animals in a frame falls below an empirical threshold (0.8 in our work) and the distance between these two animals is less than the initial animal size, it indicates that this frame (lost target frame) contains a lost target. Next, UDMT detects an abrupt jump in movement speed (based on variance and max/mean) and a sudden drop in localization confidence to determine which animal is assigned the incorrect ID through majority voting. Finally, the position of the lost animal is re-localized by detecting the animal mask (segmented by the localization refining module in the lost target frame) without ID. The trajectory of the lost target is fixed through backward tracking. Besides, if the position of a specific ID stays outside the animal mask (off-target localization) for more than 3 s, it is identified as a missing target. The same procedure will be applied to correct it.

### Automatic parameter tuning module

Tracking accuracy is highly related to search region size, which is codetermined by the search region scale and target size bias (Supplementary Table 1a). The target size is the side length of the tracking box and is defined as the initial animal size plus the target size bias. The search region scale is the scale factor of the search region size relative to the target size. The search region size is equal to the target size multiplied by the search region scale. The search region scale (1.5, 2, or 2.5) and target size bias (−10%, 0, or 10% of the initial target size) are automatically adjusted based on proposed evaluation metrics, including the number of ID corrections, off-target localizations, and missing targets. They are strongly correlated to tracking performance (Supplementary Table 1b) and can be calculated without ground truth. The number of ID corrections reflects how often the ID correction module is used and has the strongest correlation with tracking performance. In most cases, fewer ID corrections indicate that the parameters used are more suitable. The number of off-target localizations refers to the cumulative number of instances where the predicted animal position of UDMT falls outside the animal mask segmented by the localization refining module. Off-target localization lasting more than 3 s is considered to be a missing target, necessitating the ID correction module to reassign an ID to it. The number of missing targets can reflect tracking accuracy.

The detailed flowchart of automatic parameter tuning is shown in Extended Data Fig. 2. We adopted conditional iterative optimization to find the best parameters (the target size bias and search region scale) that can lead to the smallest evaluation metrics. At the beginning of the iteration, we initialized the current optimum to a very large value (1000). After processing each frame, the number of ID corrections is compared to the current optimum. If the value is larger than the current optimum, the tracking using this search region size is terminated. If the number of ID corrections equals the current optimum, further conditions are evaluated sequentially, including the number of off-target localizations and missing targets. The tracking process continues if none of these metrics is larger than the current optimum or the number of ID corrections is smaller than the current optimum. If the final frame is processed, the optimal evaluation metrics are updated. The iteration continues overall all optional values of the search region size until finished. Additionally, the processing time serves as a constraint on algorithm efficiency. If other parameters have the same values, the parameter set with the shortest processing time will be adopted. Generally, a video segment of about one minute is sufficient for automatic parameter tuning. The best parameters will be used both in training and inference.

### Ablation study

To evaluate the effectiveness of the transformer architecture and the three modules in UDMT (localization refining, ID correction, and automatic parameter tuning), we conducted ablation studies using behavioral recordings of seven mice (67 Hz frame rate, 29,550 frames, N=5 videos). DiMP^63^ was used as the baseline representing CNNs, which has a similar architecture as UDMT and substitutes the transformer blocks with convolutional blocks. The backbone network of DiMP was initialized with the open-source pretrained weights (DiMP-50) publicly available at https://github.com/visionml/pytracking/blob/master/MODEL_ZOO.md. The model was fine-tuned for 20 epochs. For the experiments removing the ID correction module, we randomly reassigned IDs to the missing target of two intersecting trajectories. Backward tracking was retained to restore missing trajectories. For those experiments without automatic parameter tuning, we selected the initial target size and typical search region scale (2.0) as the parameters to obtain representative tracking performance.

### Method comparison

We compared the performance of UDMT with three baseline methods: DLC^18^, SLEAP^19^, and IDT.ai^21^. All methods were implemented with publicly available code provided by companion papers. Human annotators were instructed to label keypoints for each dataset to generate the training dataset for supervised methods (DLC and SLEAP). For mice, eight keypoints (snout, left ear, right ear, shoulder, three spine points, and tail base) were annotated for 100 frames per video. For *Drosophila*, eight keypoints (head, left eye, right eye, thorax, left midleg, right midleg, left hindleg, and right hindleg) were annotated for 30 frames per video. For *C. elegans*, seven keypoints (head, five body points, and tail) were annotated for 50 frames per video. Annotated frames were sampled from original videos using uniform clustering. DLC models were trained for 500,000 iterations as the loss converged. For the training of SLEAP, the maximum number of instances was set to match the number of animals. The second spine point, thorax, and third body point were designated as the anchor points for mice, *Drosophila*, and *C. elegans*, respectively. For the inference of SLEAP, a flow-shift tracker was applied. The maximum number of instances and tracks were both set to the number of animals to ensure realistic tracking. The ‘connect single track breaks’ option was applied for all videos. For DLC and SLEAP, the anchor point representing the centroid of each animal was used to localize the animal. For IDT.ai., we applied background subtraction for all videos and manually adjusted intensity and area thresholds using the GUI to segment individual animals. Missing values of these methods were estimated using linear interpolation based on the nearest available data. All hyperparameters not mentioned here were set as default values.

### Locomotion analysis

To obtain the instantaneous velocity of a given frame, we computed the Euclidean distance between the animal coordinates of the current frame and the next frame, and then divided by the time interval between the two frames. To calculate acceleration, we calculated the difference in the instantaneous velocity of two adjacent frames and divided by the time interval between the two frames. To identify directional changes, we resampled the trajectory at 10-frame intervals to convert it into a path of contiguous line segments with a constant time gap. Turning angles were defined as the smallest angle between adjacent segments.

### Calcium imaging analysis

Raw calcium imaging data from the head-mounted miniaturized microscope were motion-corrected by the open-source NormCorre algorithm in non-rigid mode^64^. The data were then processed by DeepWonder^65^ to extract neuronal masks and calcium traces. DeepWonder was fine-tuned to make it suitable for miniaturized microscopes, following the guidelines provided at https://github.com/yuanlong-o/Deep_widefield_cal_inferece/tree/main/Alternative/DeepWonder. For spike inference, we used the MLspike algorithm^66^, which was ranked first in the Spikefinder challenge^67^. Before being fed into the spike inference pipeline, all calcium traces were divided by their mean values for normalization. The recommended parameters for the GCaMP6f calcium indicator were used to ensure optimal performance in spike inference.

### Neuroethological analysis

For the mouse with the miniaturized microscope, high moving speed was defined as any speed larger than the mean speed. Short distance was defined as any distance below the lower quartile of the average distance between the mouse with the miniaturized microscope and its companion, which means that they were close to each other. These events must last longer than five seconds to be recognizable. Neurons exhibiting significantly increased average activity during the high-speed movement were classified as upregulated neurons (P < 0.05, two-sided Wilcoxon test). We performed an independent sample t-test to assess whether there was a significant difference in the speed of the mouse with and without the miniaturized microscope. We also assessed whether the speed of the mouse and their distance influenced their neuronal spike rate. These variables were divided into two groups: one consisting of values greater than the upper quartile and the other consisting of values less than the lower quartile. Before the t-test, we verified the homogeneity of variance between the two groups using Levene’s test. P values are indicated with asterisks. All analyses were performed by customized MATLAB scripts.

### Performance metrics

Tracking performance is assessed with comprehensive metrics including HOTA, MOTA, IDF1, and IDSW. Among them, the dominant metric is HOTA, which is computed by the code provided at https://github.com/JonathonLuiten/TrackEval. It can balance the effect of detection and association into a unified metric for explicit comparison^35^. In terms of detection, its accuracy is quantified by the overall detection accuracy (DetA), measuring the consistency between the network prediction and corresponding ground truth using intersection over union (IoU). Since a predicted object may overlap with multiple ground-truth annotations, the Hungarian algorithm is applied for one-to-one matching. Matched pairs are true positives (TPs), while unmatched predicted detections are false positives (FPs) and unmatched ground-truth detections are false negatives (FNs). For a given IoU threshold *α*, DetA is calculated as:

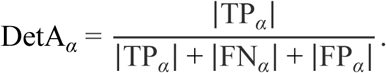

Association accuracy measures the effectiveness of a tracking model in associating detected objects to correct IDs over time. The Hungarian algorithm is used for matching and IoU is used for quantification. The intersection between two tracks is quantified by true positive associations (TPAs). Objects in the predicted track that are unmatched or matched to other ground-truth annotations are false positive associations (FPAs), while unmatched objects in the ground-truth trajectories are false negative associations (FNAs). The overall association accuracy (AssA) is calculated by averaging the Ass-IoU across all TPs in the entire dataset:

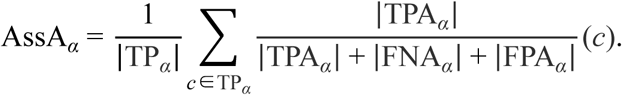

For each threshold *α*, HOTA is defined as the geometric mean of the detection and association accuracy, which ensures detection and association are evenly weighted. The final value of HOTA is obtained by integrating over different thresholds:

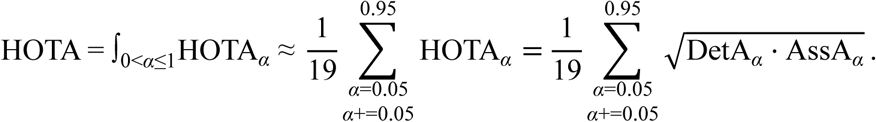

MOTA is a metric that is more sensitive to detection than association. It contains two types of detection errors (FNs and FPs) and one type of association errors (IDSW). MOTA is computed by summing these three errors, dividing by the number of ground-truth objects (gtDet), and then subtracting from one:

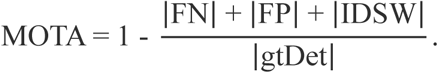

IDF1 evaluates the consistency of identity associations by balancing identity precision and recall. It is particularly effective in measuring association performance and is sensitive to ID switches, making it suitable for complex or long-duration tracking scenarios. However, IDF1 does not directly account for detection errors such as false positives and false negatives, and it needs to be complemented with other metrics like MOTA or HOTA to provide a more comprehensive evaluation of tracking performance^37^. IDF1 is formulated as:

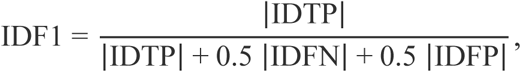

where IDTP, IDFN, and IDFP represent the number of identity true positives, identity false negatives, and identity false positives, respectively. When calculating performance metrics, the bounding box of an animal is defined as the maximum outer square centered at the position of it.

## Data availability

We have no restriction on data availability. All behavioral recordings used in this study have been archived and made publicly available at https://zenodo.org/records/14580256. The source data of neuroethological research of freely behaving mice, including behavioral recordings and synchronized calcium imaging data, are publicly available at https://zenodo.org/records/14586426.

## Code availability

All relevant resources are readily accessible on our GitHub page https://cabooster.github.io/UDMT/. The source Python code of UDMT can be found at https://github.com/cabooster/UDMT. Our specially designed user-friendly GUI and accompanying tutorial can be found at https://cabooster.github.io/UDMT/Tutorial/.

## Acknowledgements

We would like to acknowledge L. Yuan for his assistance in calcium imaging with the head-mounted miniaturized microscope. We thank G. Xiao for providing the rodents used in this study. This work was supported by the National Natural Science Foundation of China (62088102, 62222508, 62401156), Beijing Natural Science Foundation (Z240011), Beijing Laboratory of Brain and Cognitive Intelligence, Chinese Postdoctoral Foundation (2023M741962), and Tsinghua Shuimu Scholar Program (2023SM066).

## Author Contributions

Q.D., J.W. and Z. Li. supervised this research. Q.D. and X.L. conceived and initiated this project. Y.L. and X.L. designed detailed implementations, built the recording system, and performed experiments. Y.L. and X.L. developed the Python code, completed the GUI, and processed relevant data. Q.Z. and J.F. provided *C. elegans* and *Drosophila*, and assisted with relevant experiments. Y.Z., and Z. Lu provided the head-mounted miniaturized microscope and gave critical support on its imaging and data processing procedure. Y.L. and X.L. analyzed the data, prepared figures and videos, and made the companion webpage. Y. L, X.L., Q.Z., Z. Li, J.W. participated in discussions about the results. All authors participated in the drafting of the manuscript.

## Materials & Correspondence

Correspondence and requests for materials should be addressed to Q. D., J.W. and Z. Li.

---

**Extended Data Fig. 1.**
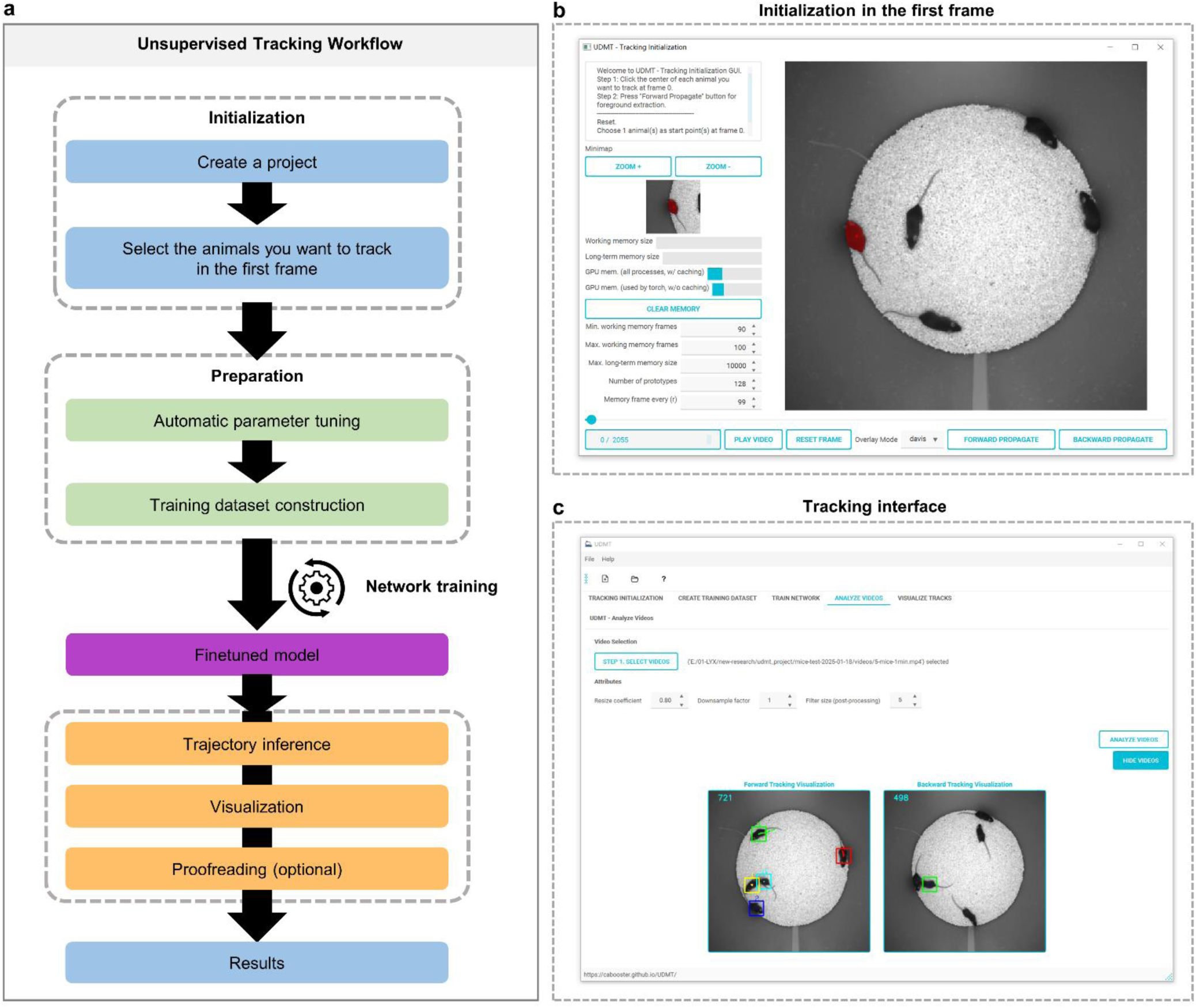
| Workflow of UDMT. **a,** Unsupervised tracking workflow. First, users just need to create a project and click the animals they want to track in the first frame. Second, parameters will be optimized and training dataset will be constructed automatically. Third, a finetuned model will be trained in an unsupervised manner. Finally, the tracking results will be obtained after network inference, visualization and proofreading. **b,** Screenshot of the interactive initialization graphical user interface (GUI). **c,** Screenshot of the tracking interface.

**Extended Data Fig. 2.**
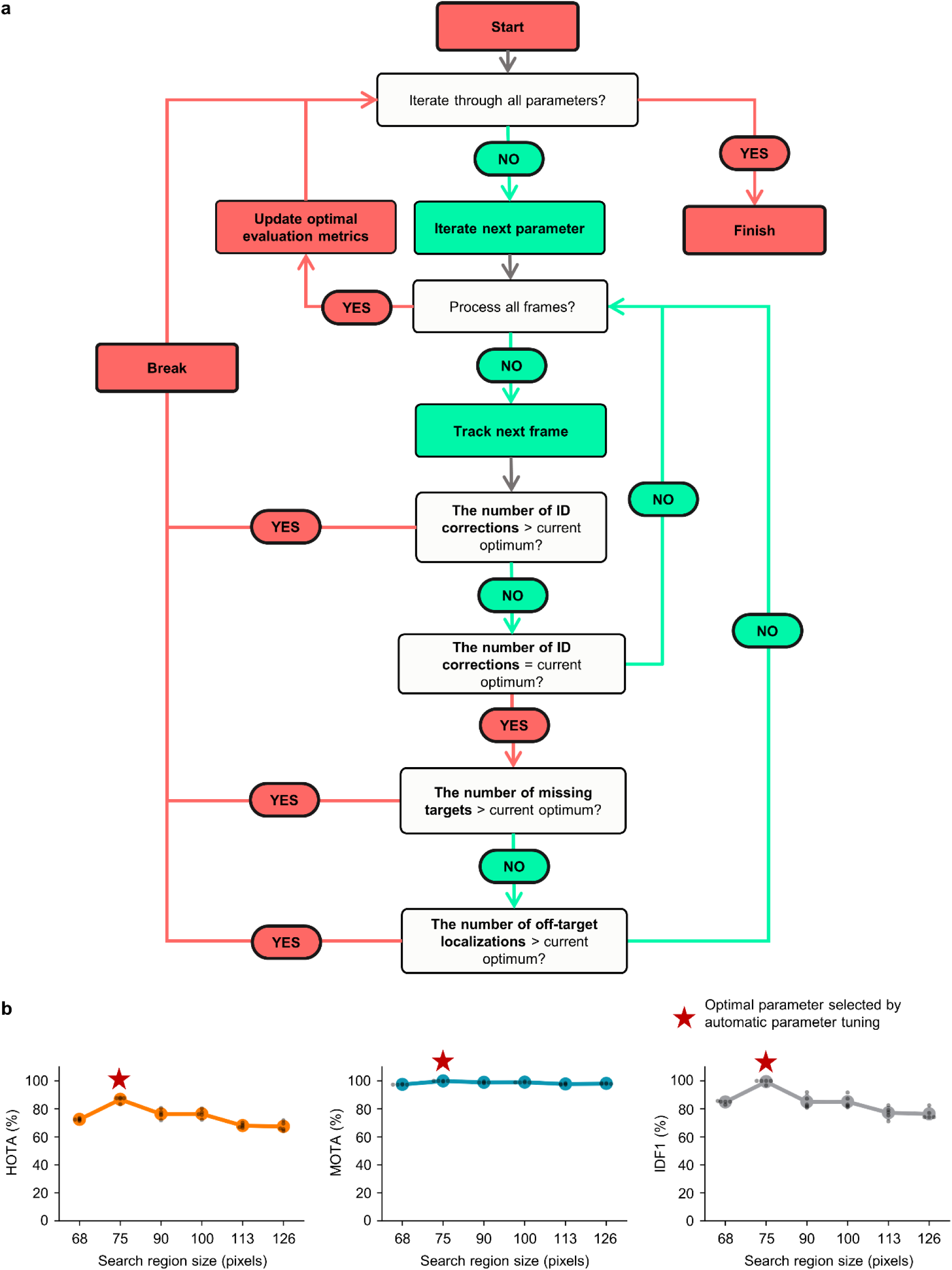
| Automatic parameter tuning module of UDMT. **a**, Flowchart of the automatic parameter tuning module. **b**, Tracking performance of UDMT with different search region sizes. The 5-mouse dataset (30 Hz frame rate, 18,000 frames) was used for quantitative evaluation. The search region size was calculated based on the search region scale and target size bias. Detailed information is provided in Supplementary Table 1. Lines represent mean values and error bars represent 95% confidence intervals. N=5 for all dataset and each gray point indicates an independent experiment.

**Extended Data Fig. 3.**
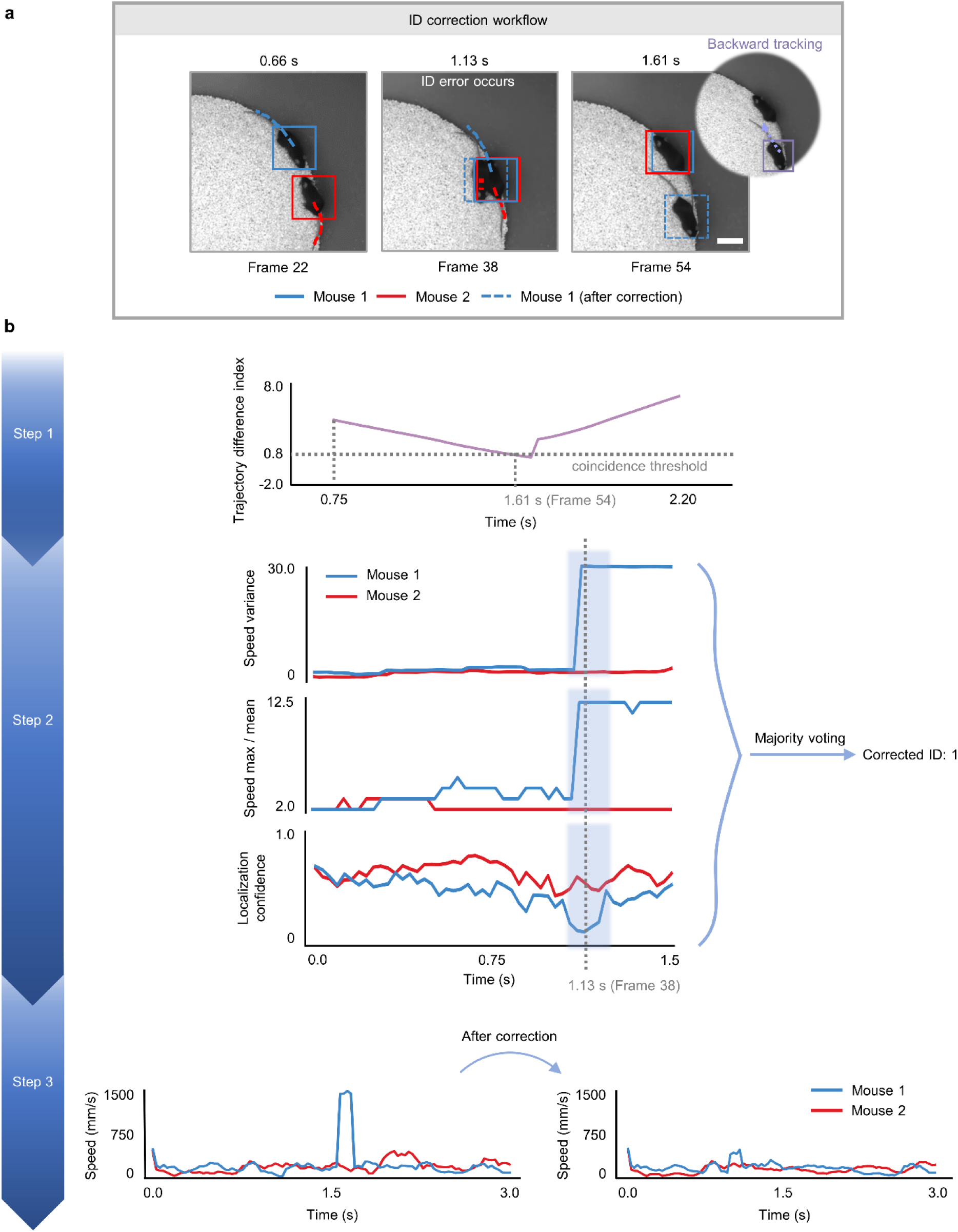
| ID correction module of UDMT. **a**, Three key frames illustrating the principle of ID correction. The lost target (mouse 1) in frame 54 is relocated and the correct ID is reassigned. Scale bar, 50 mm. **b**, Detailed ID correction workflow. First, the frame containing lost target is detected if the trajectory difference index of two mice falls below the threshold^62^. Second, UDMT detects an abrupt jump in movement speed (based on variance and max/mean) and a sudden drop in localization confidence to determine which individual (mouse 1 in this example) is assigned a wrong ID. Finally, the trajectory of the misidentified mouse was fixed using backward tracking. Line plots show the speeds of two mice as a function of time before and after ID correction.

**Extended Data Fig. 4.**
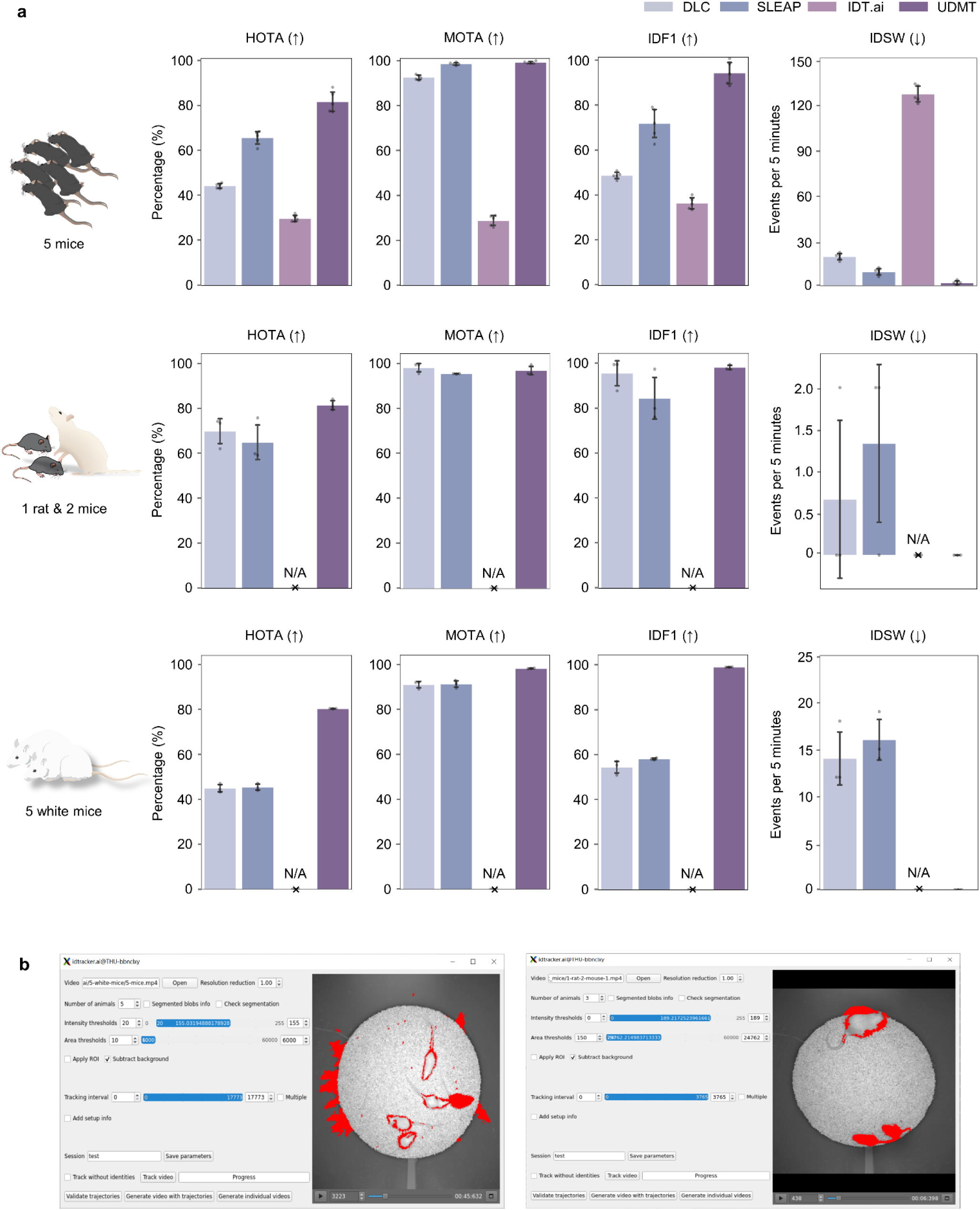
| Comparing tracking performance of various methods on rodents. **a**, Tracking performance of UDMT, DeepLabCut (DLC), idtracker.ai (IDT.ai) and SLEAP, quantified by HOTA, MOTA, IDF1, and IDSW. Bars and whiskers represent mean values and standard deviations, respectively. Each gray point indicates an independent experiment. The 7-mouse dataset (top row, 67 Hz frame rate, 29,550 frames, N=5), rat-and-mouse dataset (middle row, 68 Hz frame rate, 8630 frames, N=3) and 5-white-mouse dataset (bottom row, 70 Hz frame rate, 17,760 frames, N=3) were used for quantitative evaluation. N/A means that IDT.ai fails to track animals on the rat-and-mouse dataset and 5-white-mouse dataset. **b**, Examples showing how IDT.ai fails to segment low-contrast animals. Left, white mice. Right, white rat.

**Extended Data Fig. 5.**
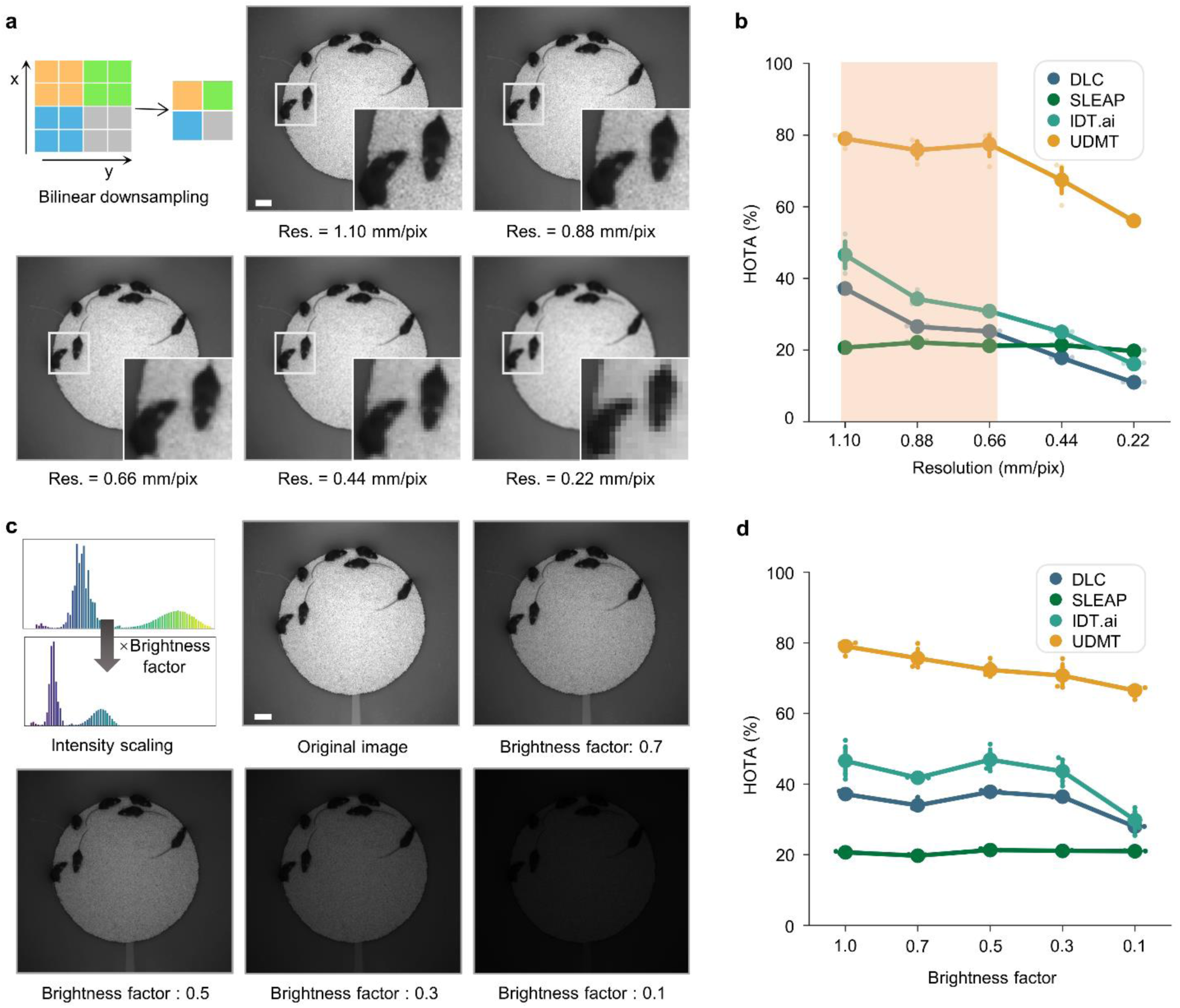
| Comparing UDMT with other methods on different recording resolutions and brightness. The 7-mouse dataset (67 Hz frame rate, 29,550 frames) was used for quantitative evaluation. **a**, Representative video frames of different resolutions. Magnified views of the boxed regions are shown at the bottom of each image. Videos of different resolutions were obtained by downsampling the original high-resolution videos with bilinear interpolation. **b**, Quantitative relationship between image resolution and tracking accuracy of different methods. The orange shaded area in the line plot indicates the range in which the performance of UDMT is robust to resolution degradation. **c**, Representative video frames under different brightness conditions. Videos of different brightness were obtained by multiplying the original bright video by different brightness factors (1.0, 0.7, 0.5, 0.3, 0.1) and rounding it into integers. **d**, Quantitative relationship between video brightness and tracking accuracy of different methods. Lines represent mean values and error bars represent 98% confidence intervals. N=5 for all datasets and each translucent point indicates an independent experiment. Scale bars, 50 mm. UDMT was finetuned with a pretrained model for 20 epochs. The results were smoothed with a five-frame window. DLC and SLEAP were implemented with the companion code of relevant papers^18,19^. For these two supervised methods, one human annotator was instructed to localize the 8 keypoints (snout, left ear, right ear, shoulder, three spine points, tail base) across 100 frames sampled from the whole video using the k-means clustering approach. All DLC models were trained for 500,000 iterations. For the training of SLEAP, the maximum number of instances was set to match the number of animals, and the second spine point was designated as the anchor part. For the inference of SLEAP, flow-shift tracking was used as the tracker method. The max instance and max number of tracks were specified as seven to ensure realistic tracking results. For these two supervised methods, the predicted second spine point representing the mouse’s body center was used to recognize the position of a mouse. All hyperparameters not mentioned here were set as default values. IDT.ai is a weakly supervised learning method and was implemented with the companion code of the relevant paper^21^. We chose reasonable parameters to segment individual animals with the GUI. All NaN values were linearly filled using the nearest non-empty result. For each method, a specified model was trained for each video resolution.

**Extended Data Fig. 6.**
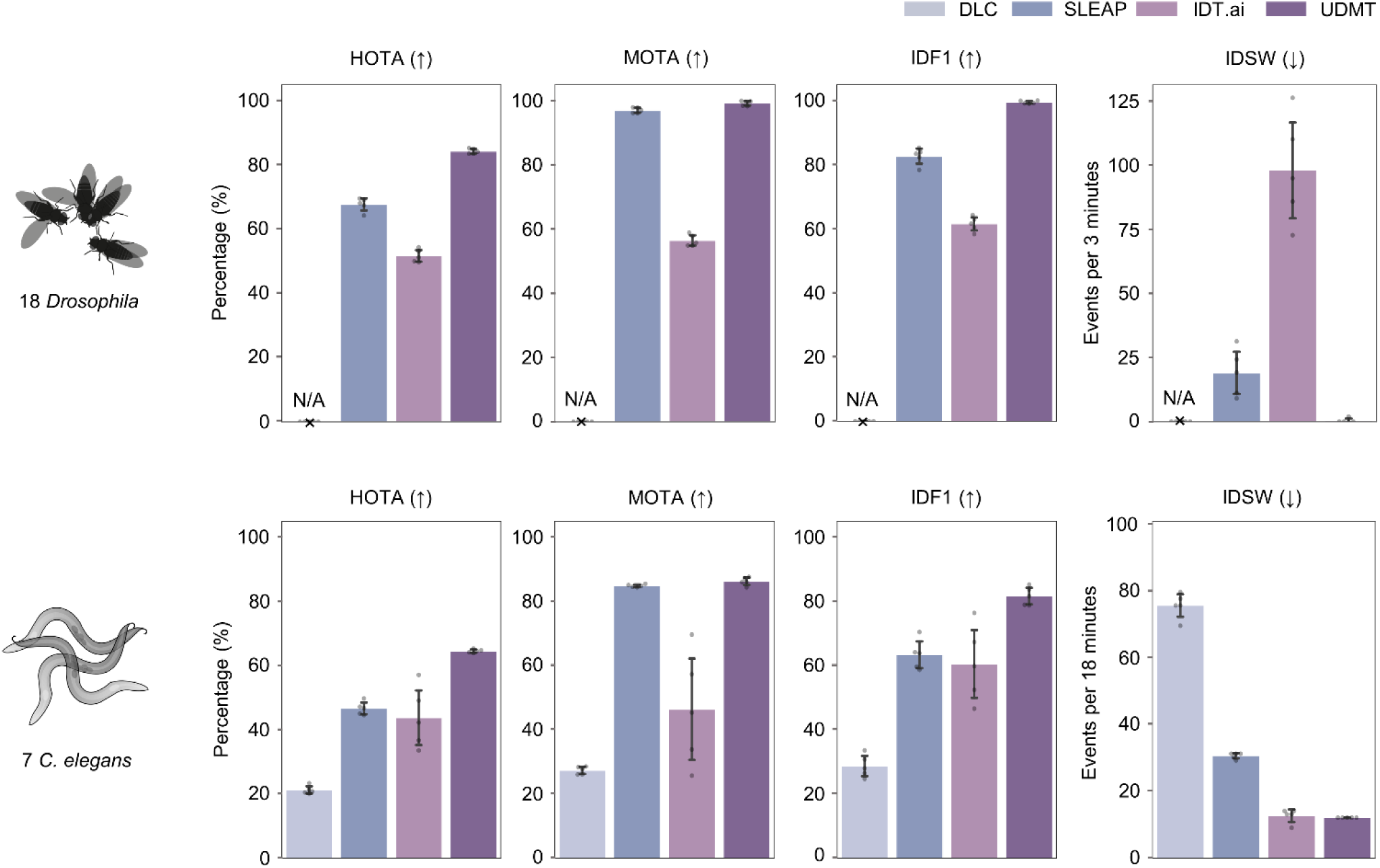
| Comparing tracking performance of various methods on *Drosophila* and *C. elegans* dataset. Tracking performance of UDMT, DLC, IDT.ai and SLEAP, quantified by HOTA, MOTA, IDF1, and IDSW. Bars and whiskers represent mean values and standard deviations, respectively. N=5 for all datasets and each gray point indicates an independent experiment. The 18-*Drosophila*-2 dataset (54 Hz frame rate, 32,370 frames) and 7-*C. elegans* dataset (10 Hz frame rate, 8,630 frames) were used for quantitative evaluation. N/A means that DLC fails to track animals on the *Drosophila* dataset.

**Extended Data Fig. 7.**
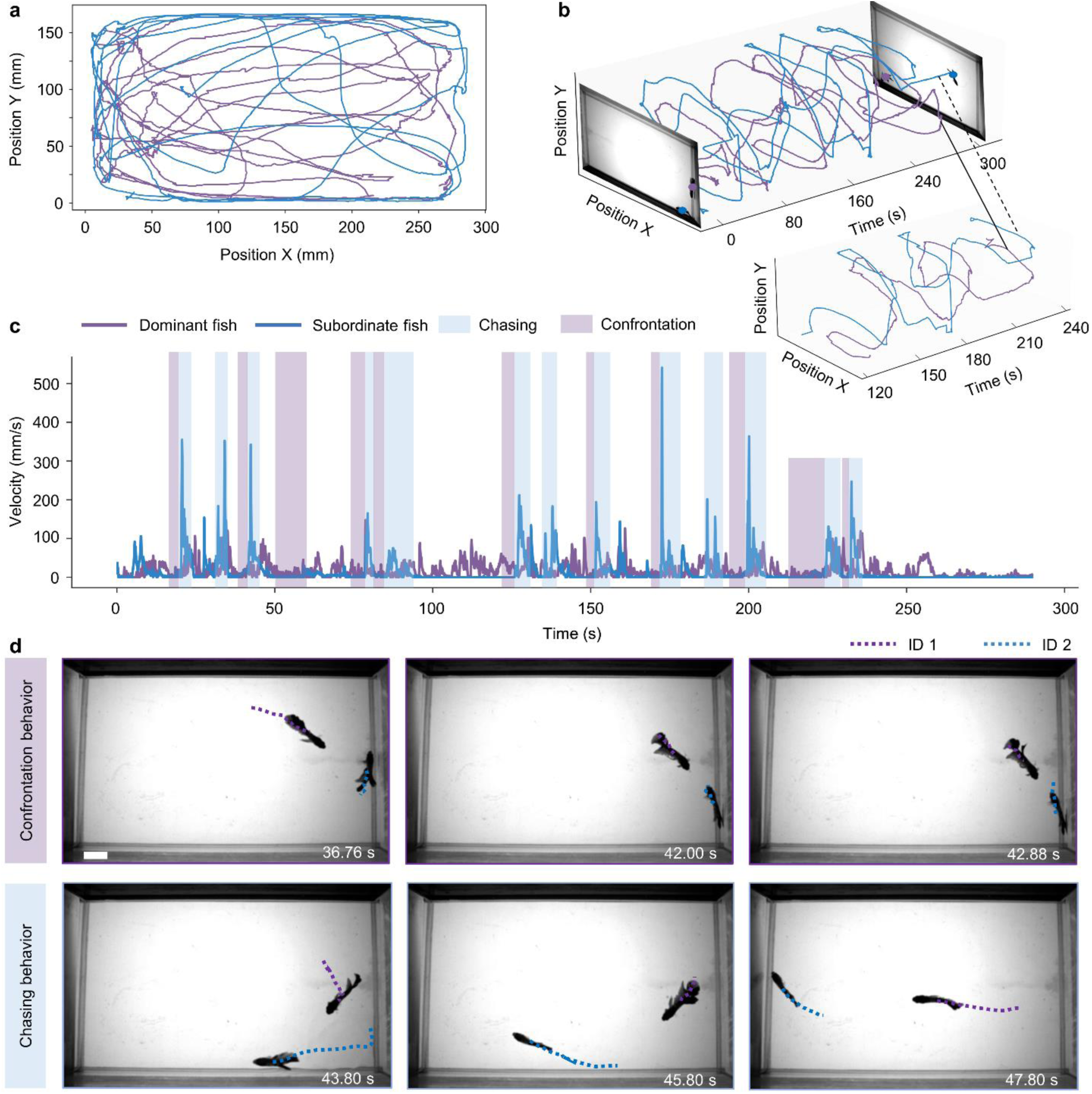
| Behavioral analysis of two *Betta splendens* using UDMT. The interaction of two *Betta splendens* (betta fish) in the same arena is continuously recorded (51 Hz frame rate, 15,020 frames). Their swimming trajectories were extracted using UDMT. **a**, Projected trajectories of the two betta fish during the entire recording period. **b**, 3D (x-y-t) trajectories of the two betta fish in a 300-second time window. The trajectories during chasing are shown separately in the inset. **c**, Velocity of the two betta fish. Traces were averaged over 10 frames. The purple shaded area indicates the time of confrontation, during which the speed of both animals decreased to nearly zero. The blue shaded area indicates the time of chasing, when the speed of the escaping fish (ID 2) increased rapidly. **d**, Representative video frames showing the confrontation and chasing behavior of the two betta fish. The tracks show the movement of the two betta fish over 1.5 seconds. Scale bar, 20 mm.

