## Supplementary Information for "Unsupervised multi-animal tracking for quantitative ethology"

### CONTENT

#### I. Supplementary Figures


| | |
|---|---|
| **Supplementary Figure 1** | Localization refining module of UDMT. |
| **Supplementary Figure 2** | Experimental setups for mouse and *Drosophila*. |
| **Supplementary Figure 3** | Tracking performance of UDMT under fluctuating illumination. Refining module of UDMT. |
| **Supplementary Figure 4** | Quantitative evaluation of UDMT at different signal-to-noise ratios (SNRs). |
| **Supplementary Figure 5** | Experimental setups for *C. elegans* and *Betta splendens*. |
| **Supplementary Figure 6** | Turning Behavior of freely moving *C. elegans*. |
| **Supplementary Figure 7** | Data dependency and training stability of UDMT. |


#### II. Supplementary Tables


| | |
|---|---|
| **Supplementary Table 1** | Detailed parameter information for the sample data in automatic parameter tuning module evaluation. |
| **Supplementary Table 2** | Summary of all datasets used. |


#### III. Supplementary Videos


| | |
|---|---|
| **Supplementary Video 1** | Tracking the movement of 10 mice simultaneously with UDMT. |
| **Supplementary Video 2** | UDMT tracks 5 white mice in low-contrast conditions. |
| **Supplementary Video 3** | Cross-species tracking with UDMT: trajectory visualization of 1 rat and 2 mice. |
| **Supplementary Video 4** | Tracking rapid movements with UDMT. |
| **Supplementary Video 5** | Neuroethology analysis of multiple mice combined with a head-mounted microscope. |
| **Supplementary Video 6** | Tracking the movement of 17 *Drosophila* simultaneously with UDMT. |
| **Supplementary Video 7** | Tracking the movement of 7 *C. elegans* simultaneously with UDMT. |
| **Supplementary Video 8** | Analyzing the aggressive behavior of betta fish with UDMT. |

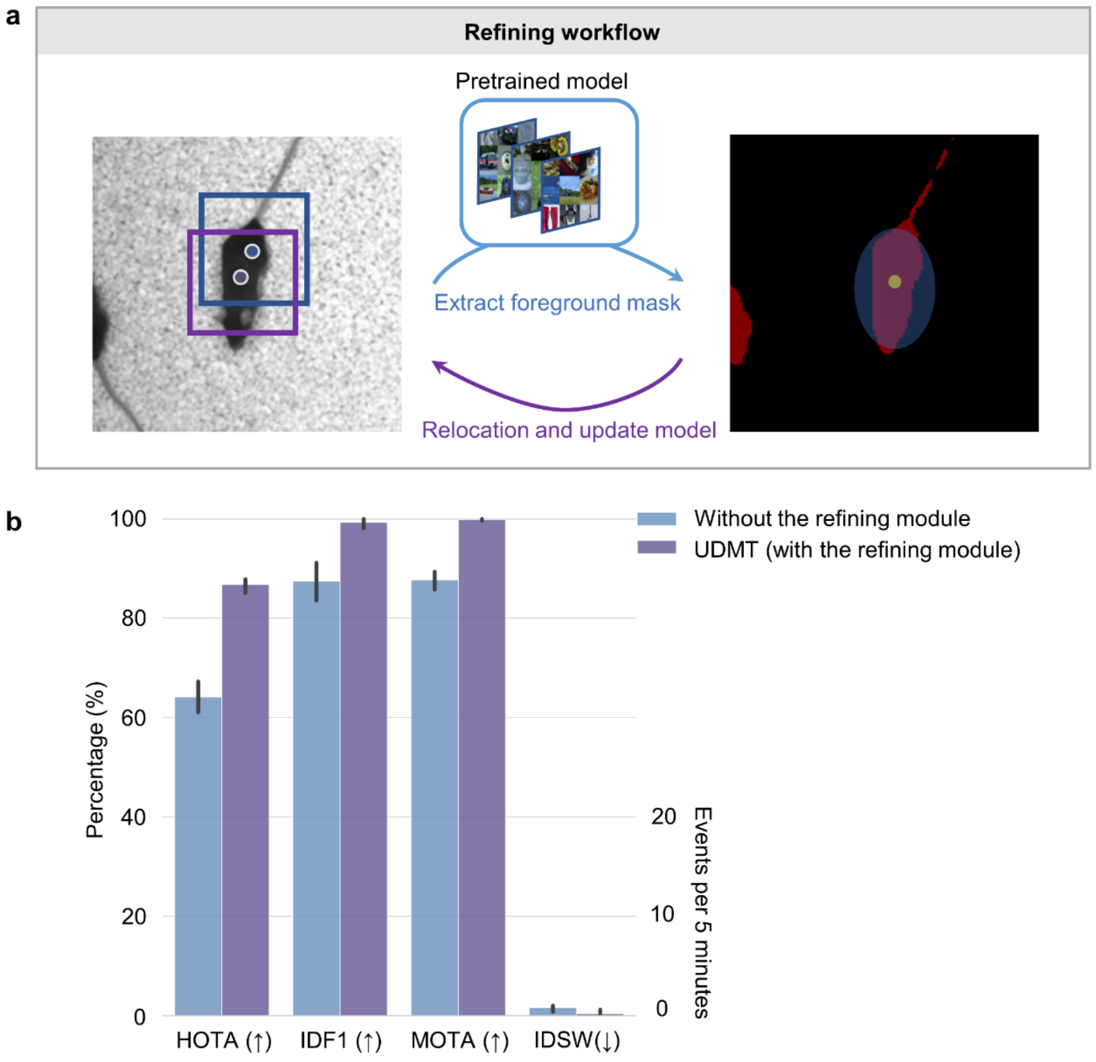


**Supplementary Figure 1**

**Localization refining module of UDMT.**

**a,** Schematic of the localization refining module. Users click the individuals they want to track to initialize the training process, and the individuals selected by users will be automatically segmented by a generalized model[1] and sent to a video object segmentation model[2] for animal segmentation throughout the entire video. This module locates the animal masks based on the initial position tracked by UDMT, and optimizes the results to the center of gravity of corresponding segment masks. Refined positions are used to update the tracking model. **b**, Tracking performance (quantified by HOTA[3], MOTA[4], IDF1[5] and ID switches) of UDMT with and without refining module. Bars represent mean values and error whiskers represent 95% confidence interval. Videos recording the movement of 5 mice (30 Hz frame rate, 18,000 frames, N=5) were used for quantitative evaluation.

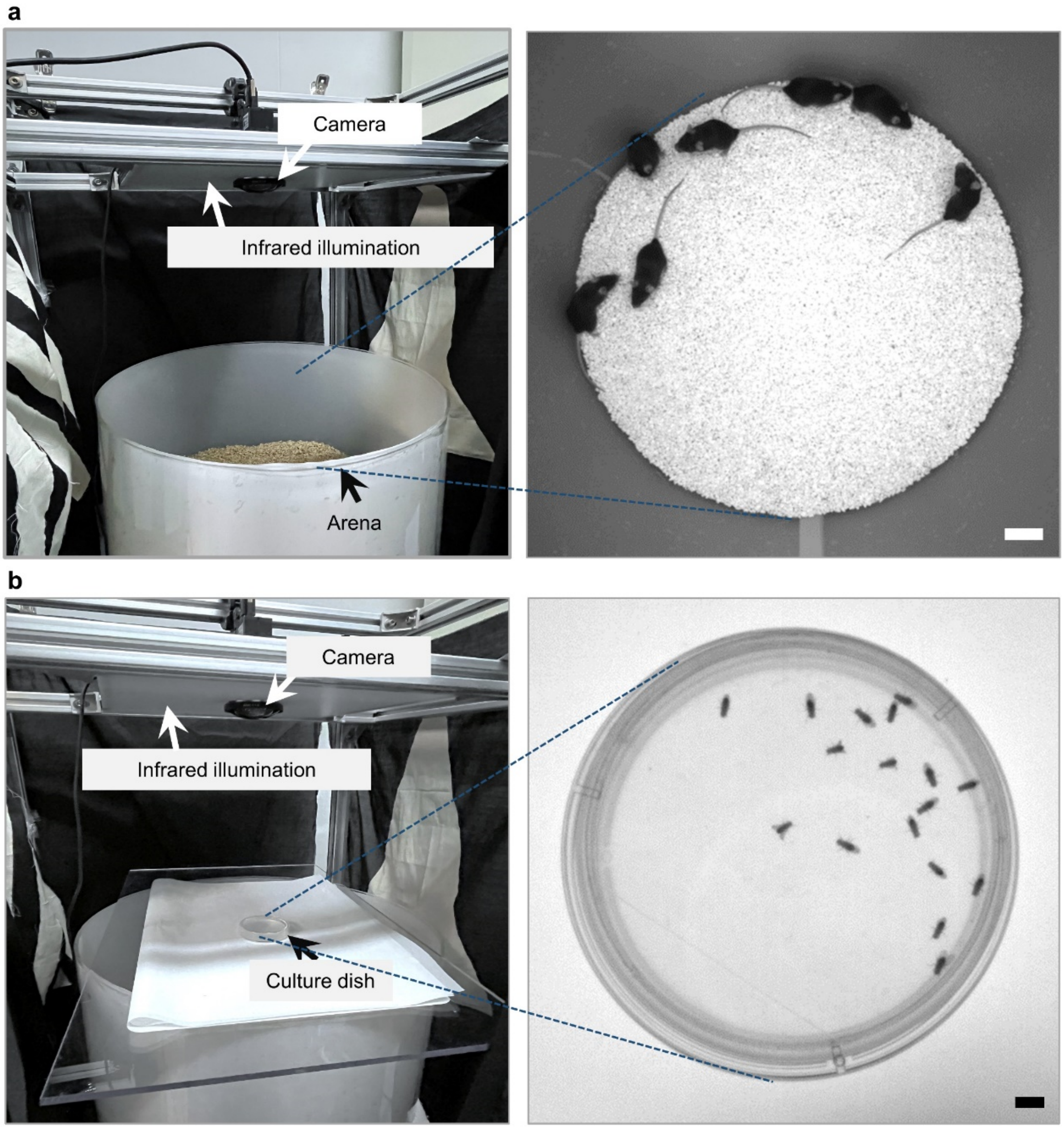


**Supplementary Figure 2**

**Experimental setups for mouse and *Drosophila*.**

**a**, Exterior view of the setup used to record mouse (left) and a representative frame from a video of 7 mice (right). Scale bar, 50 mm. **b**, Exterior view of the setup used to record *Drosophila* (left) and a representative frame from a video of 17 *Drosophila* (right). Scale bar, 5 mm.

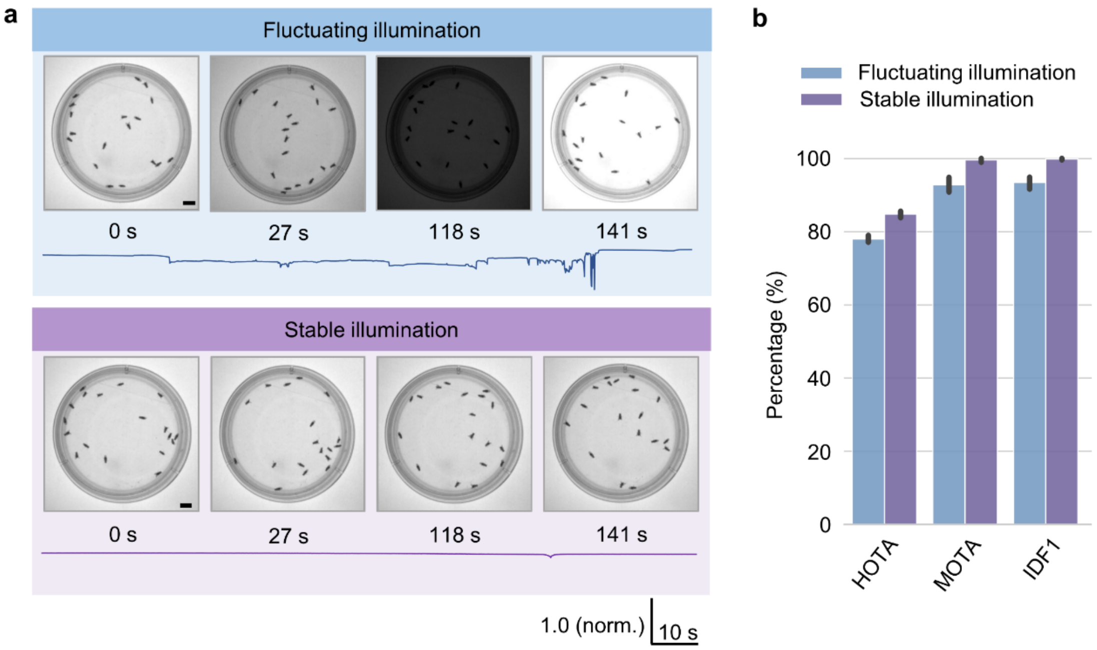


**Supplementary Figure 3**

**Tracking performance of UDMT under fluctuating illumination.**

Videos recording the movement of 18 *Drosophila* (50 Hz frame rate, 7,050 frames) were used for quantitative evaluation. Infrared illumination flickered irregularly during recording to generate illumination fluctuations. **a**, Representative video frames under fluctuating and stable illumination. Line plot under each group of images shows the change of illumination intensity over time. Scale bar, 5 mm. **b**, Tracking performance (quantified by HOTA, MOTA, and IDF1) of UDMT under fluctuating and stable illumination (N=3). The model was finetuned with a pretrained model for 20 epochs and the last epoch was used for comparison. Bars represent mean values and error whiskers represent 95% confidence intervals.

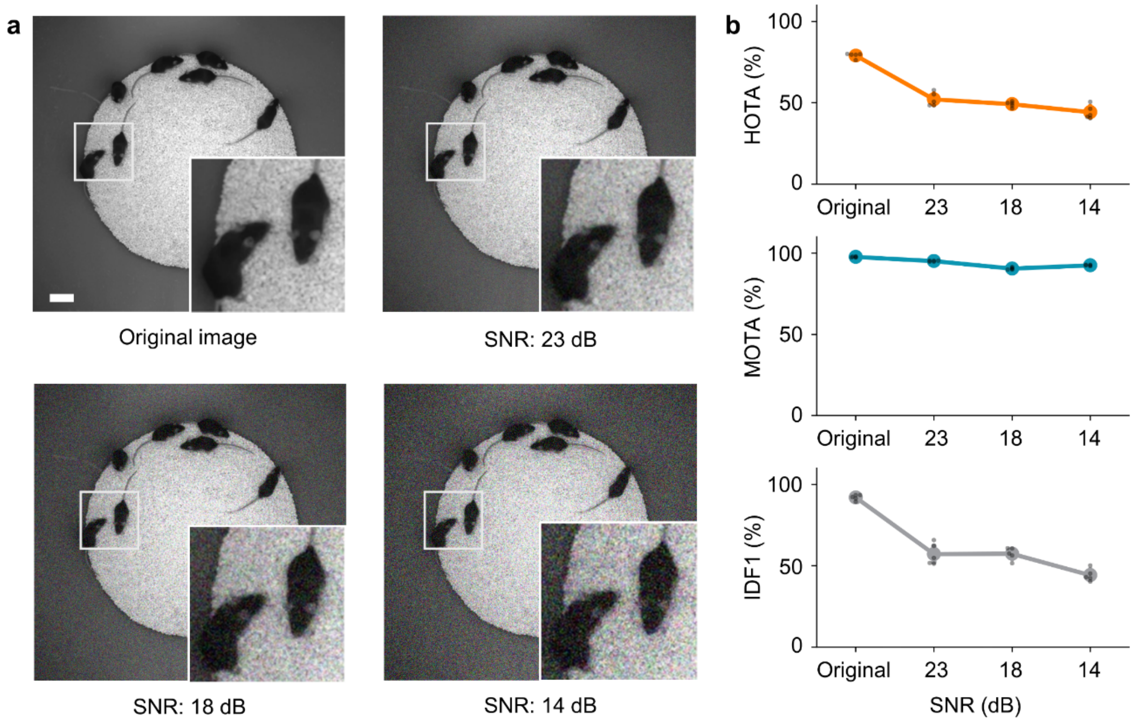


**Supplementary Figure 4**

**Quantitative evaluation of UDMT at different signal-to-noise ratios (SNRs).**

Videos recording the movement of 7 mice (60 Hz frame rate, 29,550 frames) were used for quantitative evaluation. Different SNR levels were implemented by adding Gaussian noise of different standard deviations. A specified model was trained for each SNR level. All models were finetuned with a pretrained model for 20 epochs and the last epoch was used for comparison. **a**, Representative video frames at different SNR levels. Magnified views of the boxed regions are shown at the bottom of each image. Scale bar, 50 mm. **b**. The relation between tracking performance of UDMT and the SNR of video. Lines represent mean values and error bars represent 95% confidence intervals. N=5 for all datasets and each gray point indicates an independent experiment.

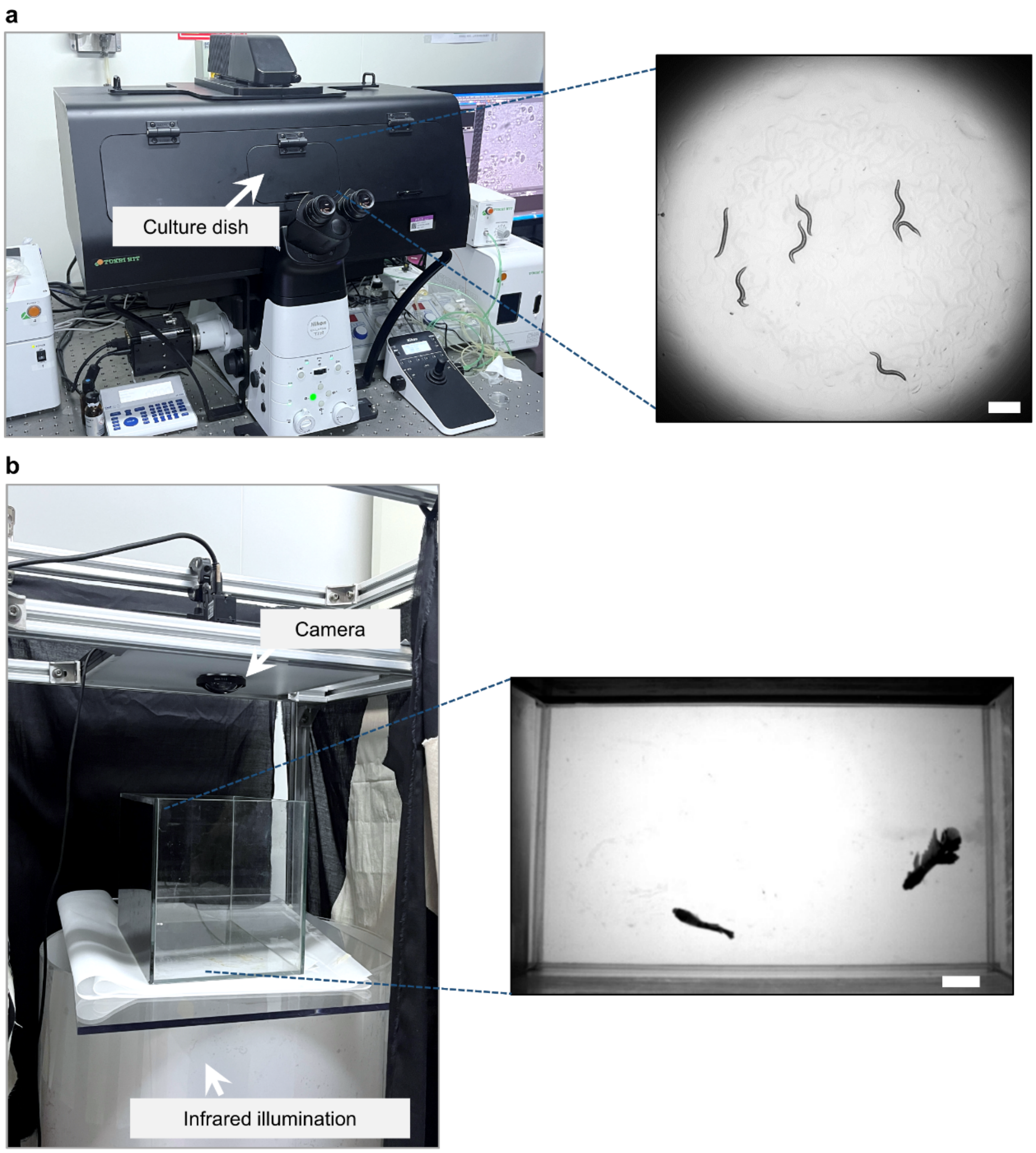


**Supplementary Figure 5**

**Experimental setups for *C. elegans* and *Betta splendens*.**

**a**, Exterior view of the setup used to record *C. elegans* (left) and a sample frame from a video of 7 *C. elegans* (right). Scale bar, 300 μm. **b**, Exterior view of the setup used to record *Betta splendens* (left) and a sample frame from a video of two *Betta splendens* (right). Scale bar, 20 mm.

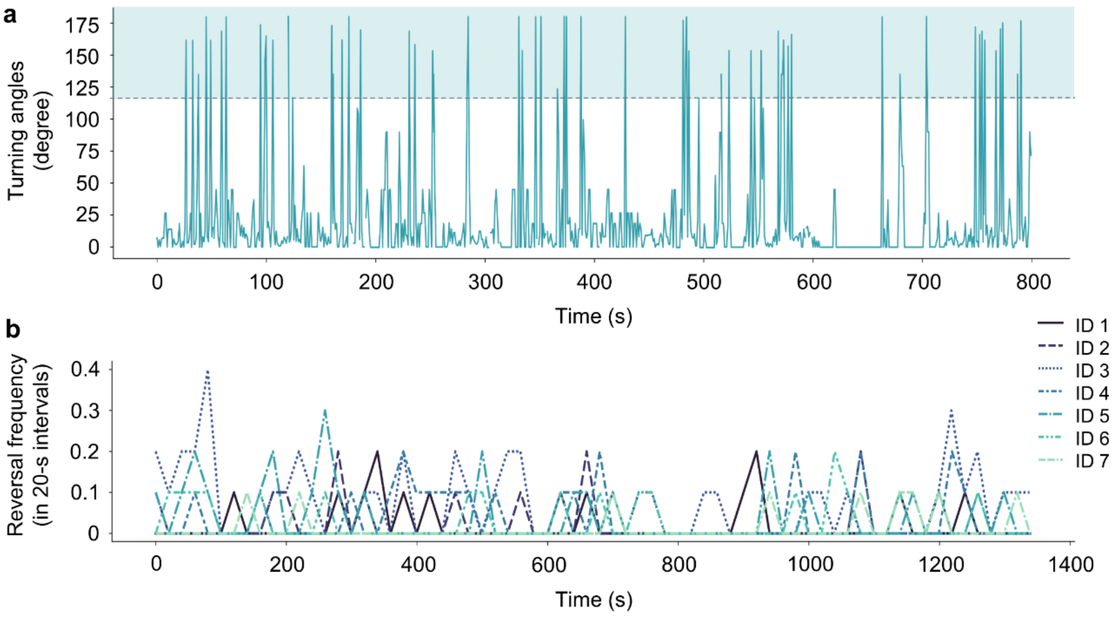


#### Supplementary Figure 6

##### Turning behavior of freely moving *C. elegans*.

Videos recording the movement of 7 *C. elegans* (10 Hz frame rate, 14,550 frames) were used for behavioral analysis. The model was finetuned with a pretrained model for 20 epochs and the last epoch was used. **a**, Turning angles of a *C. elegans* in an 800-second time window. Angles change of greater than 120 degrees (the dashed line) are defined as directional change[6]. **b**, Reversal frequency of the 7 *C. elegans*. Reversal frequency was averaged over 20-s intervals.

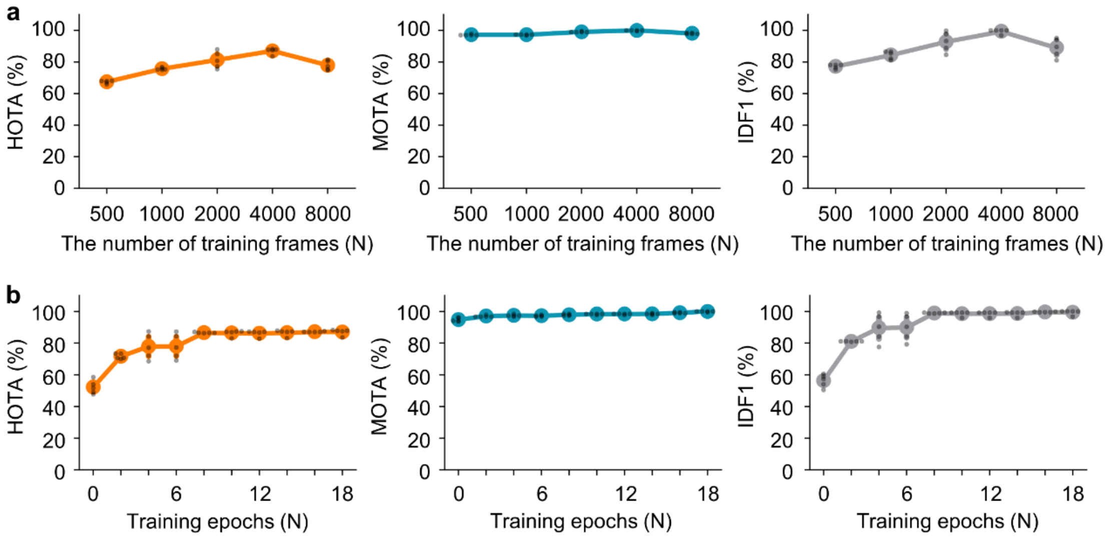


**Supplementary Figure 7**

**Data dependency and training stability of UDMT.**

Videos recording the movement of 5 mice (30 Hz frame rate, 18,000 frames) were used for quantitative evaluation. **a**, The relation between tracking performance of UDMT and the number of training frames. **b**, Tracking performance with the increase of training epoch. The first point (training epoch = 0) indicates tracking performance of the pretrained model. Lines represent mean values and error bars represent 95% confidence intervals. N=5 for all datasets and each gray point indicates an independent experiment.

**Supplementary Table 1**

**Detailed parameter information for the sample data in automatic parameter tuning module evaluation.**

**a**, The search region size was determined based on the search region scale and target size bias. **b**, The number of ID corrections, missing targets, off-target localizations and processing time used for the first 4,000 frames were calculated to select the optimal search region size. The corresponding HOTA values for the entire video are also shown. The 5-mouse dataset (30 Hz frame rate, 18,000 frames, N=5) was used for quantitative evaluation. HOTA indicates the mean value of 5 samples.

**a.**

| **Target size bias** \ **Search region scale** | **1.5** | **2** | **2.5** |
|---|---|---|---|
| **-10** | 68 | 90 | 113 |
| **-5** | 75 | 100 | 126 |

**b.**

| **Search region size (pixels)** | **68** | **75** | **90** | **100** | **113** | **126** |
|---|---|---|---|---|---|---|
| The number of ID corrections (↓) | 2 | **1** | 4 | 10 | 12 | 8 |
| The number of missing targets (↓) | 0 | **0** | 0 | 0 | 0 | 0 |
| The number of off-target localizations (↓) | 0 | **0** | 0 | 0 | 0 | 1 |
| Processing time (↓) | 436 s | **310 s** | 329 s | 377 s | 391 s | 364 s |
| HOTA (%) | 73.0 | **86.4** | 76.4 | 76.4 | 68.1 | 67.3 |

**Supplementary Table 2**

**Summary of all datasets.**

| Species | Animals | Image size | Resolution | Duration | FPS (Hz) | Labels (frames) | ID |
|---|---|---|---|---|---|---|---|
| Mouse (black) | 3 | 652 x 636 | 0.92 px/mm | 6’57” | 67 | 94 | √ |
| | 5 | 652 x 636 | | 8’04” | 66 | 107 | |
| | 5 | 416 x 444 | | 11’30” | 94 | 140 | |
| | 7 | 664 x 630 | | 7’21” | 67 | 99 | |
| | 10 | 600 x 686 | | 6’00” | 40 | 48 | |
| Mouse with miniscope | 5[a] | 640 x 512 | | 5’03” | 64 | -- | × |
| | | | | 5’36” | 64 | -- | × |
| Mouse (white) | 5 | 704 x 642 | | 4’13” | 70 | 60 | √ |
| Rat & Mouse | 3 | 664 x 630 | | 2’06” | 68 | 29 | √ |
| *Drosophila* | 12 | 560 x 624 | 6.09 px/mm | 3’36” | 55 | 80 | √ |
| | 17 | | | 8’35” | 54 | 186 | |
| | 18 | | | 4’32” | 54 | 99 | |
| | 18[b] | | | 2’10” | 54 | 47 | |
| | 23 | | | 9’57” | 54 | 216 | |
| *C. elegans* | 6 | 614 x 612 | 0.15 px/um | 24’16” | 10 | 98 | √ |
| | 7 | | | 22’35” | | | |
| *Betta splendens* | 2 | 528 x 350 | 1.56 px/mm | 4’49” | 51 | -- | × |
| | | | | 4’11” | | | |

a. one with a head-mounted microscope.

b. video with illumination change.
